# DNA virus infections shape transposable element induction *in vitro* and *in vivo*

**DOI:** 10.1101/2024.04.18.589901

**Authors:** Jiang Tan, Vedran Franke, Eva Neugebauer, Sandra Pennisi, Melanie Anna Schächtle, Justine Lagisquet, Anna Katharina Kuderna, Stephanie Walter, Isabelle Welker, Armin Ensser, Markus Landthaler, Thomas Stamminger, Thomas Gramberg, Altuna Akalin, Emanuel Wyler, Florian Full

## Abstract

Transposable elements (TEs) are implicated in a variety of processes including placental and preimplantation development and a variety of human diseases. TEs are known to be activated in the context of some viral infections, but the mechanisms and consequences are not understood. We show strong activation of TEs upon DNA virus infection, in particular the MLT- and THE1-class of LTR-containing retrotransposons as well as a subset of LINE-1-, Alu-elements and HERVs. Mechanistically, two key pathways induce TEs upregulation: inhibition of the KAP1/TRIM28 repressive complex by phosphorylation, and expression of the pioneer transcription factor double-homeobox 4 (DUX4), which is known to be involved in TE-induction during zygotic genome activation in embryonic development. DUX4 is induced by DNA viruses, it binds to TEs upon infection and analysis of genes adjacent to TEs shows pathways that are important for DNA virus infections. Analysis of knockdown, knockout and overexpression data reveal that almost all TEs expressed upon herpesviral infection are regulated by KAP1/TRIM28 and DUX4. Interestingly, analysis of single cell sequencing data from patients with DNA virus-associated cancers showed that *in vivo* TEs expression strongly correlates with virus infection, indicating a possible role in viral oncogenesis.

## Introduction

Transposable elements (TEs) are DNA segments that have the ability to move within the genome (1). TEs are reminiscent of past virus infections that resulted in integration into the host genome but lost the ability to produce virus particles and to propagate horizontally. The human genome is composed of approximately 45% TEs (2). TEs can be classified into two categories: class-I transposons (retrotransposons) and class-II transposons (DNA transposons). DNA transposons contain a transposase and have a mobile DNA intermediate, while retrotransposons require an RNA intermediate and a reverse transcriptase (3).

Retrotransposons are of scientific interest due to their use of an RNA intermediate and self-replicating life cycle for dissemination, similar to retroviruses. Retrotransposons are classified by function and structure into LINE (long interspersed nuclear elements), SINE (short interspersed nuclear elements), and LTR (long terminal repeats) (4–5). Once dismissed as parasitic or “junk” DNA, they have emerged as integral components with functional significance within host genomes. Most TEs in the human genome lost the ability to “jump” or change their position in the genome but can still be transcribed and some of them encode for proteins. Proteins originally encoded by retrotransposons have been co-opted for regulating essential processes, including placental cytotrophoblast fusion, preimplantation development, and intracellular RNA transport across neurons, a process named retrotransposon exaptation (6–8). Retrotransposon exaptation significantly expands mechanisms of gene regulation and transcript diversity of host genes by enriching them with numerous cis-regulatory elements. This is particularly true for retrotransposons harboring intact LTR elements with intrinsic promoter and enhancer activities, as well as splicing donor/acceptor sequences (9–12).

Although retrotransposons can be threats to genome integrity, and are therefore generally inactivated through degenerative mutations or epigenetic silencing, a subset of them are robustly induced and tightly regulated in specific developmental, physiological, and pathological contexts. These contexts include preimplantation development, germ cell development, immune response, aging, and cancer (13–18). Proteins like SETDB1 or SUMO-modified KAP1 (TRIM28), associated with histone methylation and Krüppel-associated box domain (KRAB) zinc finger proteins (ZNFs), have been implicated in the repression of TEs (19–25). In addition, it is known that multiple TEs are activated during zygotic genome activation (ZGA) in early embryonic development. ZGA is driven by the pioneer factor double homeobox 4 (DUX4), which is exclusively expressed during the 4-cell to 8-cell state in human embryonic development and is known to induce TEs expression (26–29). Certain viral infections, such as SARS-CoV-2, Influenza A virus, HIV-1 and DNA viruses have also been shown to induce the expression of retrotransposons, including endogenous retroviruses (ERVs) (30–33). This suggests a potential link between viral infection and retrotransposon expression. However, the molecular mechanisms of TEs-activation still remain poorly understood.

We recently discovered that DUX4 is upregulated during herpesviral infection (34), making it an excellent model for exploring the mechanisms of controlled induction of TEs. To address these gaps, our goal was to comprehensively analyze TEs-transcription upon herpesvirus infection. Additionally, we examined single cell RNA-sequencing (RNA-seq.) data of cancer patients that suffer from tumors caused by DNA viruses, namely Merkel cell polyomavirus, human papillomavirus, and Epstein-Barr virus, confirming *in vivo* the induction of TEs in viral cancers. We demonstrate that HSV-1 infection induces the phosphorylation of KAP1, which is found to bind to the DUX4 promoter, thereby modulating the expression of both DUX4 and TEs. Phosphorylation of KAP1 has been shown to regulate sumoylation and KAP1 repression negatively (35, 36). This implies that viral infection leads to TEs expression through regulation of KAP1 and induction of DUX4. This investigation explores the complex relationship between DNA viruses, germline transcription factor DUX4, and TEs and provides new insights into the regulatory networks that control TEs activation.

## Materials and Methods

### Cell culture

HEK 293T, human foreskin fibroblasts (HFF) and SLK endothelial cells were cultured in Dulbecco’s modified Eagle’s medium (DMEM, Thermo Fisher Scientific), supplemented with 10% (vol/vol) heat-inactivated fetal bovine serum (FBS, Thermo Fisher Scientific), 2 mM GlutaMAX (Thermo Fisher Scientific), 1 mM HEPES (Thermo Fisher Scientific), and 1% penicillin-streptomycin (vol/vol). The maintenance medium for DUX4-inducible cells consisted of DMEM supplemented with 10% (vol/vol) FBS, 2 mM GlutaMAX (Thermo Fisher Scientific), 10 mM HEPES buffer (Thermo Fisher Scientific), 1% penicillin-streptomycin (vol/vol), and 1 µg/ml Puromycin. DUX4 overexpression was initiated by the addition of 0.2 µg/ml doxycycline to the culture medium. KSHV lytic replication in iSLK.219 cells (37) was induced by the addition of 1 µg/ml doxycycline to the culture medium. All cell lines were cultured under standard conditions, and monitoring for mycoplasma contamination was performed monthly using the MycoAlert Kit from Lonza.

### Generation of shKAP1 HFF cell lines and DUX4 inducible 293T cell lines

For the Generation of shKAP1 HFF cell lines, 5 x 10^6^ HFF cells were seeded in 10-cm dishes in Dulbecco’s minimal essential medium (DMEM) (Gibco) supplemented with 10 % fetal calf serum (FCS, Sigma-Aldrich), and 1 % penicillin-streptomycin (PS, Sigma-Aldrich). Utilizing the Lipofectamine 2000 reagent (Invitrogen), the cells were co-transfected on the next day with packaging plasmids pLP-VSVg (38) and pCMVdeltaR8.9 (kindly provided by T. Gramberg) together either with the plasmid pAPM-D4-miR30-L1221 (a gift from Jeremy Luban; Addgene #115846) (39) expressing a control shRNA or plasmid pAPM-D4 miR30-TRIM28 ts3 (Addgene #115864) (39) encoding for a shRNA targeting KAP1. 48 hours post transfection, the supernatant was harvested, filtered through a 0.45-μm filter and used for transduction of HFF cells. This was done by seeding HFF in 6-wells (8 x 10^4^/well) and incubating them for 24 h with 50 µl of the regarding lentiviral supernatant and 7.5 μg/mL polybrene (Sigma-Aldrich). To select for successfully transduced HFF, the cells were then provided with medium (DMEM, 10 % FCS, PS) containing 5 µg/ml puromycin. The knockdown of KAP1 was verified via western blotting and immunofluorescence. For the Generation of DUX4 inducible 293T cell lines, the processes are similar to above. Transfections were carried out using GenJet (SignaGen Laboratories) or Lipofectamine 2000 (Thermo Fisher Scientific) following the respective manufacturer’s protocols. The DUX4-inducible plasmid pCW57.1-DUX4-CA was obtained from Addgene (#99281).

### Western Blotting

Cells were lysed using RIPA HS buffer (10 mM Tris-HCl pH 8.0, 1 mM EDTA, 500 mM NaCl, 1% Triton X-100 (vol/vol), 0.1% SDS (vol/vol), 0.1% deoxycholic acid (DOC)), supplemented with Aprotinin and Leupeptin, MG-132, and sodium metavanadate (Sigma-Aldrich). The cell pellet was subjected to centrifugation at 4°C, 14,000 rpm for 30 minutes. Subsequently, the resulting samples were diluted with a Laemmli-SDS sample buffer and heated for 5 minutes at 95°C.

### RNA-seq. library preparation

HFF/shCtrl and HFF/shKAP1 were seeded in duplicates in 6-wells (3 x 10^5^/well) in medium without selection (DMEM + 10 % FSC + PS). The day after, the medium was harvested and stored at 37 °C. The cells were either mock infected (1.5 ml fresh medium) or infected with the HCMV laboratory strain TB40/E wild type (40) (1.5 ml medium/virus mix) to obtain a multiplicity of infection (MOI) of 1. The viral titer was defined based on immediate early (IE) protein-forming units as described elsewhere (41). 2 hours post infection (hpi) the cells were provided with additionally 1.5 ml of the previously conditioned medium. 48 hpi the cells were harvested with 700 µl TRIzol Reagent (Invitrogen) per well and stored at -80°C.

HAP1 DUX4 ko cells were seeded in 6-well plates and infected with HSV-1 GFP (MOI 1) in PBS supplemented with 0.1% Glucose and 1% FCS. After 1 h at 37 °C the remaining virus was washed away with a low pH buffer (40 mM Citric acid, 10 mM KCl, 135 mM NaCl, pH3). The cells were harvested at 4,6,8 and 10 hpi with 700 µl TRIzol Reagent (Invitrogen) per well and stored at -80°C.

For the library of DUX4 inducible HFF and 293T cell lines, DUX4 overexpression was initiated by the addition of 0.2 µg/ml doxycycline for 8 and 24 h, 8 and 12h respectively. For the library of HCMV (BAC2) and KSHV infected cells, HFF cells were mock infected or infected with HCMV (BAC2) for 12, 36 and 72h, and SLK endothelial cells were mock infected or KSHV induced for 24, 48 and 72h. RNA was isolated using the Direct-zol RNA Miniprep Kit (Zymo Research) and further subjected to rRNA depletion using the riboPOOL kit (Biozym). To clean up the RNA in the process of rRNA depletion, AMPure beads XP (Beckman Coulter) were utilized. Afterwards, 5 ng of the resulting RNA was used for library preparation using the NEBNext® Ultra™ II Directional RNA Library Prep Kit from Illumina® (New Englands Biolabs).

### CUT&RUN

Primary HFF cells were infected with HSV-1 GFP (MOI of 1) in PBS supplemented with 0.1% Glucose and 1% FCS. The remaining virus was washed away with a low pH buffer (40mM Citric acid, 10mM KCl, 135mM NaCl, pH3) after one hour at 37°C. CUT&RUN was performed using the option 1 of the CUT&RUN protocol (v3, dx.doi.org/10.17504/protocols.io.zcpf2vn). 1µg anti-KAP1, SETDB1, H3K9me3 and H3K14ac (Abcam) or 1µg rabbit IgG (Cell Signaling Technology) were used accordingly and 5pg Drosophila spike in DNA was added to each sample for normalization. Libraries were generated using the NEBNext Ultra II DNA Library Prep Kit.

### RT-qPCR of TEs

A549 cells with KAP1-wt (KAP1-knockout rescued with wild-type KAP1), KAP1-S824A (KAP1-knockout rescue with reconstitution of KAP1 S824A) and KAP1-S473A (KAP1-knockout rescue with KAP1 S473A) (kindly provided by Benjamin G. Hale) were infected with HSV-1 for 8 hours, TRIzol reagent (Invitrogen, Carlsbad, CA, USA) was added immediately after collection, and stored at -80°C until RNA extraction followed by purification with the RNeasy Mini Kit (Qiagen, Hilden, Germany), with the homogenization step extended to 5 minutes. Genomic DNA was removed by on-column DNase I treatment (Qiagen) for 15 minutes at room temperature, and absence of contamination was confirmed by no-RT control PCR (Cq > 35). RNA from three independent experiments was quantified using a Qubit 4 Fluorometer (Thermo Fisher Scientific, Waltham, MA, USA) with the Qubit RNA HS Assay Kit, and integrity was assessed with an Agilent 2100 Bioanalyzer RNA 6000 Nano Kit (Agilent Technologies, Santa Clara, CA, USA), yielding a RIN of 8.5 and a GAPDH 3’/5’ Cq ratio of 1.2; inhibition was ruled out by serial RNA dilutions (1:10, 1:100) showing a ΔCq of ∼3.3. One-step quantitative RT-qPCR targeting TEs was performed using 50 ng total RNA in 20 µL reactions with the Universal One-Step RT-qPCR Kit (New England BioLabs, E3005L; containing Taq polymerase at 0.05 U/µL and Luna Reverse Transcriptase) and SYBR Green I, following the supplier’s protocol (reverse transcription at 55°C for 10 minutes, initial denaturation at 95°C for 2 minutes, then 40 cycles of 95°C for 15 seconds and 60°C for 1 minute), run on an Applied Biosystems QuantStudio 3 (Thermo Fisher Scientific). Specificity was confirmed by a single melt curve peak at 82°C, no-template controls (NTCs) showed Cq = 38, and standard curves had a slope of -3.35, y-intercept of 35.2, r² of 0.995, and 98.5% efficiency (10^(-1/slope) - 1), with Cq variation of 0.25 at 10 copies/µL. Primers (synthesized by Integrated DNA Technologies, Coralville, IA, USA) included GAPDH (NM_002046.7, 120 bp amplicon) for normalization [ACAGTCAGCCGCATCTTCTT (forward), TTGATTTTGGAGGGATCTCG (reverse)] and published TE-specific primers (42) MER11A [AATACACCCTGGTCTCCTGC (forward), AACAGGACAAGGGCAAAAGC (reverse)], LTR10A [ACAACTTTCCCACCAGTCCT (forward), GCAGGAGTATGAGCCAGAGT (reverse)], THE1C-int [CCAACCCGACATTTCCCTTC (forward), AGGGGCCAAGGTACATTTCA (reverse)], AluSx3 [TGAGGTGGGCTGATCATGAG (forward), TGCAACCTCCACCTCCTAAG (reverse)], AluJr [AGGCTGAGTTGGGAGGATTG (forward), GCAGGATCTCATTCTGTTGCC (reverse)], and MIRb [AGTGCCAGCTTTGGTTTCAG (forward), AGACGAGAACACTGAGGCTC

(reverse)].

### Genes and TEs expression analysis in bulk RNA-seq

For bulk RNA-seq. analysis we employed the snakePipes mRNA-seq. pipeline to process the sequencing data (43). Adapters and low-quality bases were removed using TrimGalore (https://github.com/FelixKrueger/TrimGalore). The resulting trimmed reads were aligned to the human genome (hg38) using the STAR aligner package (44). Subsequently, featureCounts (45) was utilized to count the aligned reads, generating a count matrix for downstream analyses. To visualize the coverage, bigwig files were generated through deepTools bamCoverage (46), employing size factors calculated using DESeq2 (47). For analysis of TEs expression, the snakePipes noncoding-RNA-seq. pipeline (43) was employed. Similar to gene expression analysis, preprocessing steps included the removal of adapters and low-quality bases using TrimGalore. The trimmed reads were aligned to the Repeatmasker repeat annotation, obtained from the UCSC database, utilizing the STAR aligner (44). For analysis of TEs expression in different parts of the genome (TEs located inside of genes, 5kb downstream of genes, and outside of genes), 3 corresponding TEs annotation files were made by overlapping a repeat masker file with the hg38 gene annotation file for corresponding position. To quantify TEs expression, TEtranscripts (48) was employed to count aligned TEs reads. Subsequently, the expression values were normalized and the differential expression of TEs were analyzed using DESeq2.

### Genes and TEs expression analysis in single cell RNA-seq

For the analysis of single-cell RNA-seq data obtained from patients, we employed the STARsolo pipeline to align sequencing reads to the human reference genome (hg38) (49) and corresponding viral genomes, including EBV (NC_007605), HPV (NC_001526.4), and MCPyV (NC_010277). The generated feature-barcode matrices from all individual cells were aggregated and transformed into a Seurat object using the R package Seurat (50). For the analysis of TEs at the single-cell level, we utilized the scTE package (51). TEs expression matrices from individual cells were aggregated and transformed into a Seurat object using the same Seurat package (50). We then merged the gene expression matrices and TEs expression matrices. This integrated dataset was subjected to log-normalization and linear regression using the NormalizeData and ScaleData functions of the Seurat package. Principal component analysis (PCA) was then performed, and the first 15 principal components (PCs) were utilized for Unifold Manifold Approximation and Projection (UMAP) non-linear dimensionality reduction. This was followed by the calculation of a k-nearest neighbor graph and Louvain clustering. Marker genes per cluster were identified using the FindAllMarkers function from Seurat package (50) based on the Wilcoxon Rank Sum test. To visualize gene expression patterns, the DoHeatmap function from Seurat package (50) was employed to generate heatmaps for the top variable genes, based on scaled expression values. Marker gene expression and SingleR analysis (52) were used for annotating cell-type-specific clusters. Clusters with concurrent annotations were identified and merged to generate the cell-type annotated object. For additional visualization, FeaturePlot, DotPlot and Vlnplot functions from Seurat package (50) were utilized to explore genes and TEs expression. Statistical significance was determined using two-sided nonparametric Wilcoxon rank-sum tests (*P < 0.05, **P < 0.01, ***P < 0.001, ****P < 0.0001).

### ChIP-seq analysis

For ChIP-seq analysis, we processed the data using the snakePipes DNA-mapping and the ChIP–seq pipelines. Adapters and low-quality bases were then removed employing TrimGalore (https://github.com/FelixKrueger/TrimGalore). The resulting trimmed reads underwent alignment to the human genome (hg38) using Bowtie2 (53). Reads mapping to blacklisted regions from the Encode Consortium69 were excluded from further analysis. Additionally, Picard MarkDuplicates (v.1.65; https://broadinstitute.github.io/picard/) was utilized to identify and remove duplicated reads. For subsequent peak calling analysis, only properly paired mapped reads and reads with a mapping quality over 3 were retained. Peak calling was executed using MACS2, employing the input as a control for comparison (54). To assess log2 (FC) on peak regions between mock and HSV-1 infection, CSAW (55) was employed. To visualize the ChIP-seq data, Bigwig files were generated using deepTools bamCoverage (46). To identify peaks overlapping with TEs, the obtained peaks were intersected with the Repeatmasker repeat annotation downloaded from the UCSC database.

### CUT&RUN analysis

For CUT&RUN analysis, we processed the data using the snakePipes DNA-mapping and the ChIP–seq pipelines with CUT&RUN parameters. Adapters and low-quality bases were removed by TrimGalore (https://github.com/FelixKrueger/TrimGalore). The resulting trimmed reads underwent alignment to the human genome (T2T) and the spike-in dm6 genome using Bowtie2 (53). Reads mapping to blacklisted regions from the Encode Consortium69 were excluded from further analysis. Additionally, Picard MarkDuplicates (v.1.65; https://broadinstitute.github.io/picard/) was utilized to identify and remove duplicated reads. The spike-in dm6 genome was used to calculate sizeFactors for the normalization. To visualize the CUT&RUN data, Bigwig files were generated using deepTools bamCoverage (46).

### ATAC-seq analysis

For ATAC-seq analysis, data processing was conducted using the snakePipes DNA-mapping and the ATAC–seq pipelines (43), with the DNA-mapping segment aligning data similar to the CHIP–seq analysis (see above). To ensure data quality, TrimGalore (https://github.com/FelixKrueger/TrimGalore) was employed to remove adapters and low-quality bases. The resulting trimmed reads were aligned to the human genome (hg38) using Bowtie2 (53). Reads mapping to blacklisted regions from the Encode Consortium69 were excluded from subsequent analysis. Additionally, duplicated reads were identified and filtered out using Picard MarkDuplicates (v.1.65; https://broadinstitute.github.io/picard/). For the ATAC-seq analysis, BAM files were further filtered to include only properly paired reads with appropriate fragment sizes (<150 bases). To identify accessible chromatin regions, peak calling was performed using MACS2 (54). CSAW (55) was also applied to calculate the log2 (FC) on peak regions between mock and HSV-1 infection. For data visualization, Bigwig files were generated using deepTools bamCoverage (46). To investigate peaks overlapping with TEs, the obtained peaks were intersected with the Repeatmasker repeat annotation downloaded from the UCSC database.

### Evaluation of intron and exon expression

To check the ratio of introns / exons, the intron and exon expression was evaluated by the R package INSPEcT (56) by using the corresponding BAM files from the analysis of virus infection RNA-seq. data above.

## Data visualization

The representation of overlapping genes and TEs across diverse samples was visually conveyed through Venn diagrams, generated using the R package VennDiagram (57). Expression patterns of DUX4 target genes were depicted via heatmaps created with the R-package pheatmap (https://davetang.github.io/muse/pheatmap.html). Binding patterns of DUX4 were depicted via a pieplot created with the R package webr (https://github.com/cardiomoon/webr/tree/master/R). Differential expression of genes and TEs was visually summarized through violin plots, donut plots, and volcano plots, all generated using the R package ggplot2 (58). Metagene profiles and heatmaps for both ATAC–seq and ChIP–seq data were constructed using deepTools’ plotProfile and plotHeatmap (46). While representative tracks were produced utilizing pyGenomeTracks (59). To gain insights into the functional implications, KEGG pathways enrichment analysis was performed using ClusterProfiler (60).

### Statistical analysis

P-values were computed utilizing an unpaired Student’s t-test, or one-way Anova test (Dunnett’s multiple comparison test) with statistical significance defined as P < 0.05. The padj (P-values adjusted) used in RNAseq. data analysis were adjusted P-values by using the Benjamini-Hochberg procedure to control the false discovery rate (FDR).

## Results

### TEs are strongly induced following DNA viruses infection

In order to analyze expression of TEs in response to herpesvirus infection, we generated RNA-seq. data derived from HFF cells upon HCMV (both BAC2 and TB40/E strains) infection and iSLK cells upon KSHV reactivation. In addition, we conducted an in-depth analysis of published RNA-seq. datasets derived from HFF cells both pre- and post-exposure to HSV-1 for 2, 4, 6 and 8 hours. We also performed RNA-seq. of HCMV infected HFF cells (BAC2 for 12, 36, 72 hours and TB40/E for 48 hours), and RNA-seq. datasets derived from iSLK cells both pre- and post-reactivation of KSHV for 24, 48, 72 hours. Moreover, we reanalyzed published RNA-seq. datasets derived from pre- and post-exposure to KSHV for 6, 12, 24 hours, and published RNA-seq. datasets derived from B cells pre- and post-exposure to EBV for 48 hours. These different virus infections showed high infection efficiency with increased virus load during the infection (Fig.S1A). We observed significant upregulation of numerous TEs upon herpesviral infection (Fig.1A,B,C,D; Fig.S2A,B), including the MLT- and THE-families of retrotransposons (Fig.S3A,B; Fig.S4A,B). TE-activation became evident between 2 and 4 hours post-infection (hpi) with HSV-1. Notably, from 6 hpi onwards, the upregulated TEs surged dramatically, with about 1000 TEs significantly upregulated at 8 hpi (Fig.1A). TEs of all subclasses were upregulated in the course of infection with the numbers at 8 hpi distributed as follows: 208 DNA-elements, 157 LINE-elements, 58 SINE-elements and 532 LTR-elements (Fig.S5A,B). In the context of HCMV (BAC2) infection, from 12 hpi onwards, the upregulated TEs surged dramatically, with about 948 TEs significantly upregulated at 72 hpi (Fig.1B). 786 upregulated TEs were induced by HCMV (TB40/E) at 48 hpi (Fig.1E; Fig.S2A). HCMV replication is much slower than HSV-1 replication, therefore differences in kinetics were expected. In the context of KSHV reactivation, TEs-activation became evident between 6 and 12 hours post-induction. From 24 hpi onwards, the upregulated TEs surged dramatically (Fig.1C,D), and remarkably, 92.2% of these upregulated TEs at 48hpi (654 TEs) exhibited overlap with the upregulated TEs identified in HSV-1 infection at 8hpi and HCMV (TB40/E) at 48 hpi (Fig.1E). This observed convergence underscores shared regulatory mechanisms in the TEs response to herpesviral infection. To further evaluate the impact of virus induced gene upregulation on TEs-expression, both the exon/intron ratio as well as readthrough was analyzed during virus infection. A slightly reduced exon/intron ratio was observed upon HSV-1, HCMV and KSHV infection compared to mock (Fig.S6). We then analyzed the expression of upregulated TEs that are located within genes (“in”), 5 kb downstream of genes as a marker for viral readthrough that has been observed for HSV-1 infection (“down 5kb”) (61) and outside of genes (excluding genes and 5kb downstream of genes, “out”) post virus infection compared to mock. In the context of HSV-1 infection, we observed that most of the upregulated TEs are located outside of genes, and only some of the upregulated TEs are 5kb downstream of genes in areas that are affected by HSV-1 readthrough. Very few of the TEs are located within gene bodies (Fig.S7A). Similar results were observed in the context of HCMV (Fig.S7B) and KSHV (Fig.S8) infection. Most of the upregulated TEs are located outside of genes and few of them 5kb downstream or within genes. These results indicate that the upregulation of TEs upon DNA virus infection is due to the induction of TEs located outside of genes and not just a byproduct of virus-induced gene activation or viral readthrough.

**Figure 1:**
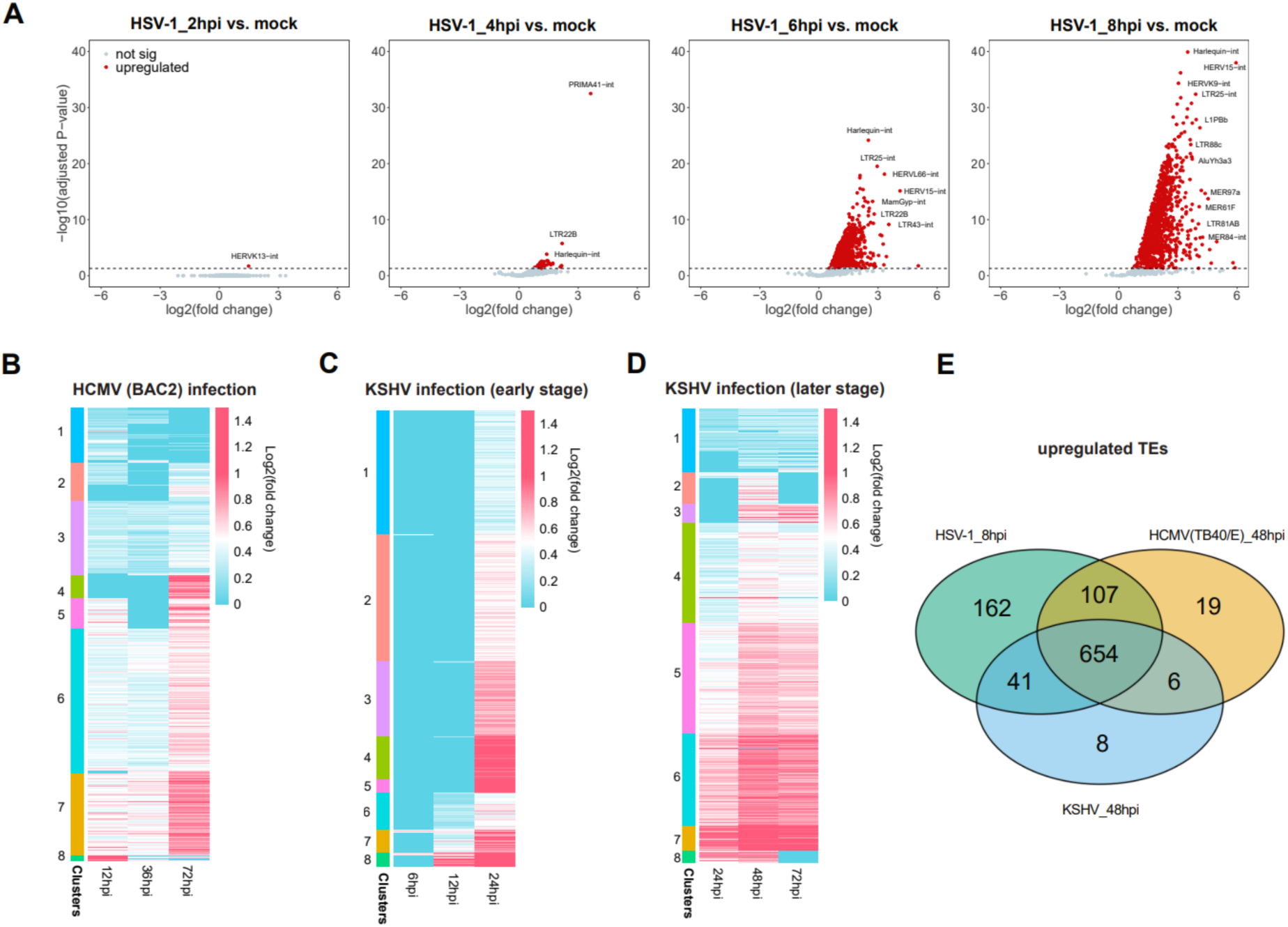
Comprehensive Analysis of TEs in Viral Infections. (**A**) Volcano plots showing significantly upregulated TEs upon HSV-1 infection of HFF cells at 2, 4, 6, and 8 hpi. Red dots indicate TEs meeting the criteria of log2(FC) > 0 and padj < 0.05. (**B**) Heatmap depicting the log2(FC) of upregulated TEs at 12, 36, and 72 hours post HCMV (BAC2) infection (hpi) of HFF cells compared to mock that cluster in 8 clusters. Values exceeding log2(FC) > 1.5 are capped at 1.5. (**C**) Heatmap depicting the log2(FC) of upregulated TEs at 6, 12, and 24 hours post KSHV infection (hpi) of SLK endothelial cells compared to mock that cluster in 8 clusters. Values exceeding log2(FC) > 1.5 are capped at 1.5. (**D**) Heatmap depicting the log2(FC) of upregulated TEs at 24, 48, and 72 hours post KSHV infection (hpi) of SLK endothelial cells compared to mock that cluster in 8 clusters. Values exceeding log2(FC) > 1.5 are capped at 1.5. Upregulated TEs meeting the criteria of log2(FC) > 0 and padj < 0.05. (**E**) Venn diagram showing the overlap of upregulated TEs induced by HSV-1 in HFF cells at 8 hpi and HCMV (TB40/E) at 48 hpi, and KSHV in SLK endothelial cells at 48hpi, providing insights into shared regulatory elements across different viral infections.

### TEs contribute to dynamic chromatin regions in response to HSV-1 infection

Next, we analyzed chromatin accessibility and histone modification patterns during HSV-1 infection to investigate the epigenetic regulation of TEs in response to herpesvirus infection. Employing transposase-accessible chromatin sequencing (ATAC-seq.) and chromatin immunoprecipitation sequencing (ChIP-seq.) targeting the transcriptional repression marks H3K9me3 and H3K27me3, we found dynamic changes in the epigenetic landscape. In uninfected cells, approximately 100,000 peaks were identified through ATAC-seq. Following HSV-1 infection, the number of peaks exhibited a declining trend over time, reaching around 60,000 peaks at 8 hpi (Fig.S9A). Notably, 18% - 23% of these peaks were associated with TEs, a proportion that decreased post-infection (Fig.S9A). To further investigate TEs with altered accessibility in response to HSV-1 infection, we compared ATAC-seq. peaks in infected and uninfected samples. The analysis revealed approximately 3,000, 12,000, and 9,000 peaks, marking sites with enhanced accessibility at 4, 6, and 8 hpi, respectively (Fig.2A,B). At 8 hpi, these upregulated ATAC peaks in repeats were distributed across 437 TEs families, with 98.4% of them exhibiting an upregulated TEs expression (Fig.2C,D).

**Figure 2:**
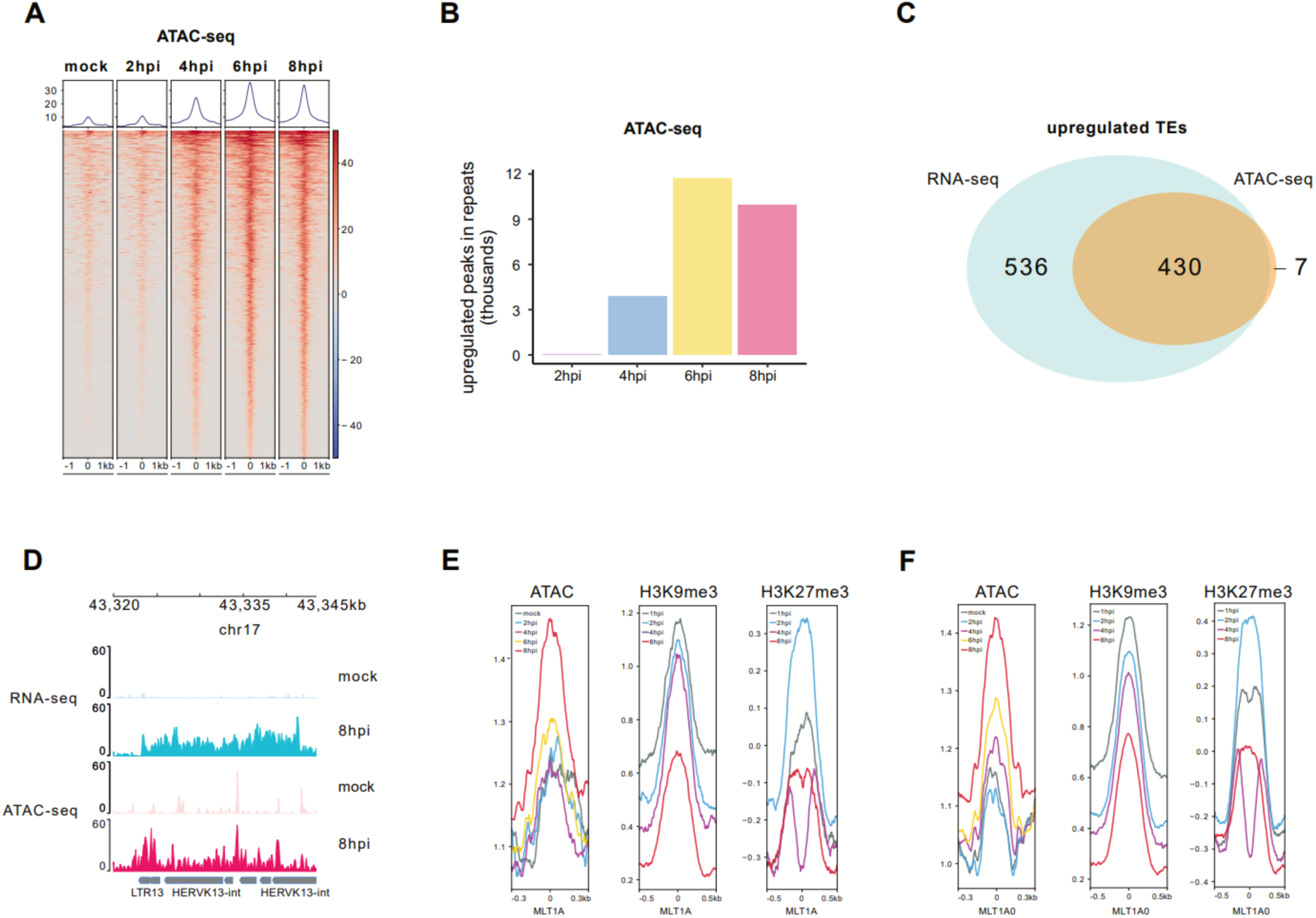
Comprehensive Analysis of Chromatin Accessibility Dynamics during HSV-1 Infection. (**A**) Heatmap displaying enrichment levels of TEs with enhanced accessibility in ATAC-seq. upon HSV-1 infection of HFF cells at mock, 2, 4, 6, and 8 hours post-infection (hpi). (**B**) Number of upregulated ATAC-seq. peaks in repeats detected in HSV-1-infected samples (HSV-1 vs. mock). (**C**) Venn diagram illustrating the overlap of upregulated ATAC-seq. peaks in 437 TEs families and upregulated TEs expression induced by HSV-1 at 8hpi. (**D**) Example genomic view illustrating TEs with upregulated expression and ATAC-seq. peaks at 8hpi. (**E**) Peak count frequency of ATAC-seq, ChIP-seq. of transcriptional repression marks (H3K9me3 and H3K27me3) peaks overlapping with MLT1A. (**F**) Peak count frequency of ATAC-seq. and ChIP-seq. of transcriptional repression marks (H3K9me3 and H3K27me3) peaks overlapping with MLT1A0.

Considering that H3K9me3 and H3K27me3 are associated with transcriptional repression, we compared ChIP-seq. peaks for these histone markers during the later stages of HSV-1 infection versus 1 hpi (62). Notably, a substantial number of H3K9me3 and H3K27me3 markers exhibited downregulation at 8hpi (Fig.S9B). For instance, TEs families like the retrotransposons MLT1A and MLT1A0 showed these distinct epigenetic patterns (Fig.2E, F; Fig.S9C). Taken together, our results show that HSV-1 infection regulates the epigenetic landscape of the human genome, particularly within TEs. The observed changes in chromatin accessibility and histone modifications at TEs suggest a potential regulatory role for these elements in the host response to viral infection.

### The pioneer transcription factor DUX4 binds to TEs upon HSV-1 infection

In a previous publication, we identified the germline transcription factor DUX4 as being induced by DNA viruses and crucial for herpesvirus infection (Fig.S10A, B, C, D, E) (34). DUX4 is known to play a role in the activation of TEs during ZGA. We therefore sought to determine whether DUX4 binding to TEs regulates their expression and that of nearby genes both in the context of HSV-1 infection and ectopic overexpression. Upon reanalyzing DUX4 ChIP-seq. data from both HSV-1-infected cells and DUX4-induced cells, we observed that 69% and 62% of the binding peaks, respectively, were located within TEs (Fig.3A,B,C). A substantial portion of these DUX4-binding TEs belonged to the MLT and THE families, with 1709 DUX4 binding sites at TEs overlapping between HSV-1-infected cells and DUX4-induced cells (Fig.3D). Further exploration involved the analysis of ATAC-seq. and ChIP-seq. data for the transcriptional repression marks H3K9me3 and H3K27me3 following HSV-1 infection. Remarkably, within the DUX4-binding TEs, we observed heightened accessibility, while H3K9me3 and H3K27me3 peaks were diminished, particularly at 8 hpi (Fig.3E; Fig.S9C). These findings underscore a dynamic interplay between DUX4, TEs, and histone modifications in response to HSV-1 infection. TEs are known to act as cis-regulatory elements regulating gene expression. Therefore, we examined the genes proximal to DUX4-binding sites within TEs and noted a significant enrichment of genes in virus-related pathways, including those associated with KSHV, HPV, HCMV, and HSV-1 infection (Fig.3F, Fig.S9D), indicating a role of DUX4 binding and TEs expression in virus infection.

**Figure 3:**
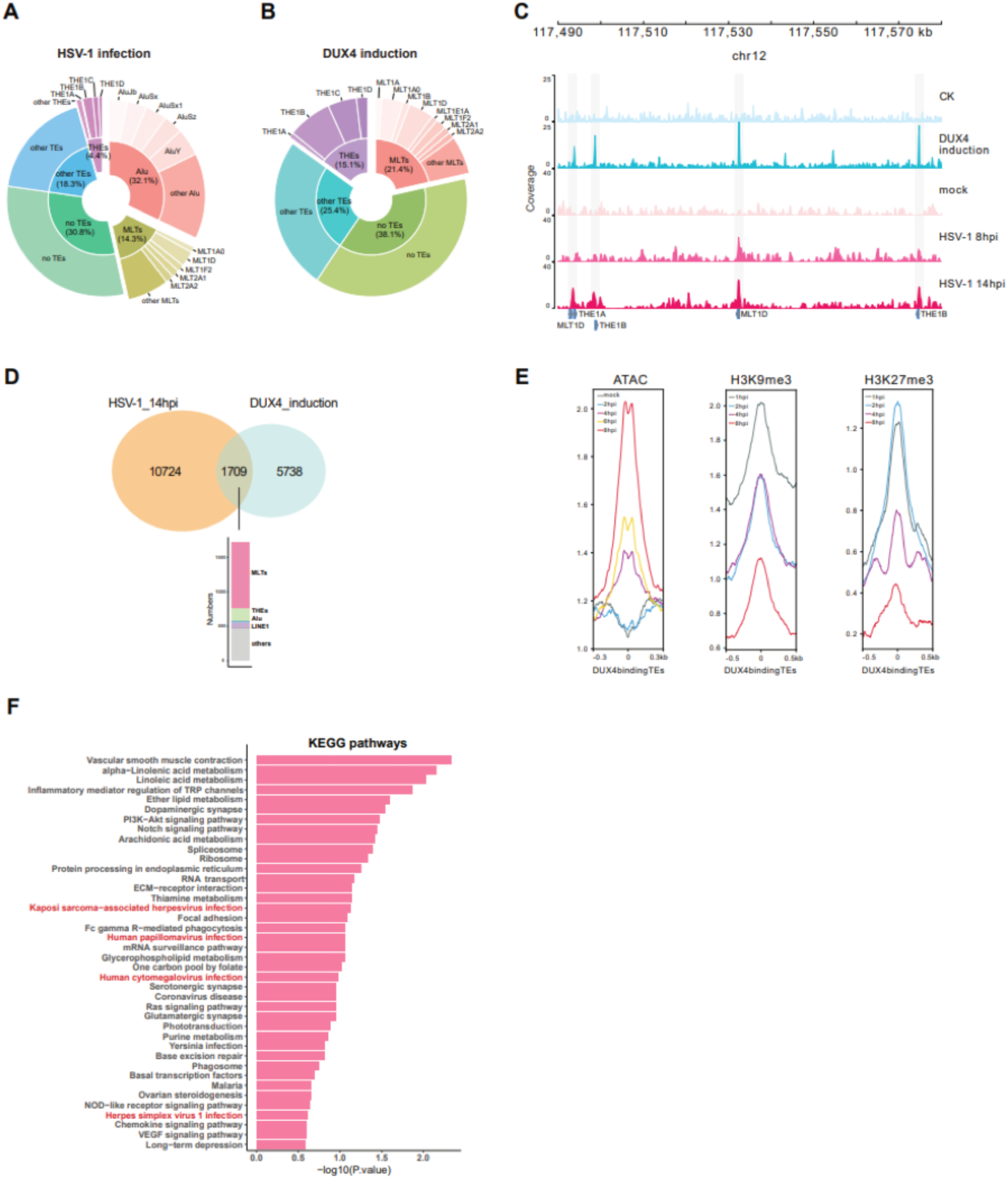
Exploring DUX4-Mediated TEs Regulation in Viral Infections. (**A**) Distribution of DUX4 binding to TEs in ChIP-seq. data of HSV-1-infected cells, categorized into THEs, MLTs, Alu, other TEs, and no-TEs. (**B**) Distribution of DUX4 binding to TEs in ChIP-seq. data of DUX4-induced HFF cells, categorized into THEs, MLTs, other TEs, and no-TEs. (**C**) RNA-Seq. and DUX4 ChIP-seq. tracks of TE binding in mock and DUX4 induced cells, as well as cells infected with HSV-1 (mock, 8 hpi, and 14 hpi). (**D**) Venn diagram illustrating the overlap of TEs bound by DUX4 at 14h post HSV-1 infection in HFF cells and DUX4 binding after DUX4 induction in HFF cells. The overlapping TEs include MLT-, THE-, Alu-, and LINE1-elements. (**E**) Peak count frequency of ATAC-seq. and ChIP-seq. of histone modification (H3K9me3 and H3K27me3) peaks overlapping with DUX4 binding at TEs. (**F**) KEGG pathways analysis of genes proximal to DUX4-binding sites within TEs (1 kb up- and downstream).

### Role of DUX4 in TEs activation

Given that approximately 70% of DUX4 binding sites are within TEs, our investigation aimed to elucidate the influence of DUX4 on TEs activation. We reanalyzed RNA-seq. datasets derived from wild-type-(wt) and DUX4 knockout-(ko) cells (Fig.S11A) at various time points (4, 6, 8, and 10 h) after exposure to HSV-1 (34). As anticipated, TEs activation surged from 6 hpi onward upon HSV-1 infection. However, upon infection of DUX4 ko cells, the expression of TEs was drastically reduced, although TEs-activation was still detectable (Fig.4A; Fig.S12), including the MLT- and THE-families (Fig.S13A,B). We observed that some activated TEs in wt cells are not regulated by DUX4 as shown by comparison of DUX4-wt and -ko cells (clusters 1, 3 and 12, Fig.4A). More than 75% of TEs demonstrated lower expression in DUX4 ko cells across all time points, with expression of TEs reduced about 99% at 10 hpi (Fig.S12A,B). The difference between TEs at 10hpi is statistically significant between DUX4-wt and -ko cells upon HSV-1 infection (Fig.S12C). Although the ko cells have a significantly lower expression of viral transcripts at 8h and 10h post infection (Fig.S1B), the big difference in TEs expression levels is unlikely to be a result of the reduced viral replication in the ko cells. Moreover, when we overexpressed DUX4 in HFF cells at 8 and 24 hours and 293T cells at 8 and 12 hours, the TEs, including MLT-, THE-, LINE-, Alu-elements and HERV-L were activated dramatically (Fig.4B,C; Fig.S14A; Supplementary Table). 448 of these upregulated TEs in DUX4 induced for 24 hours in HFF cells exhibited overlap with the upregulated TEs identified in HSV-1 infection at 10hpi in wt compared to DUX4 ko cells (Fig.S12D). Moreover, we conducted an in-depth analysis of published RNA-seq. datasets derived from DUX4-inducible myoblast cells treated with a p300/CBP inhibitor for 12 hours. p300/CBP is a histone acetyltranferase which interacts with DUX4 and is important for expression of DUX4 target genes. We observed significant upregulation of numerous TEs upon DUX4 induction, while the upregulated TEs were reduced after the addition of the p300/CBP inhibitor, including the MLT- and THE-families of retrotransposons (Fig.4D; Fig.S14B). The impact of DUX4 in the TEs expression was further confirmed by Luciferase assays (Fig.S15), demonstrating that expression of respective TEs is indeed dependent on DUX4. These findings underscore the pivotal role of DUX4 in the regulation of TEs-expression upon HSV-1 infection.

**Figure 4:**
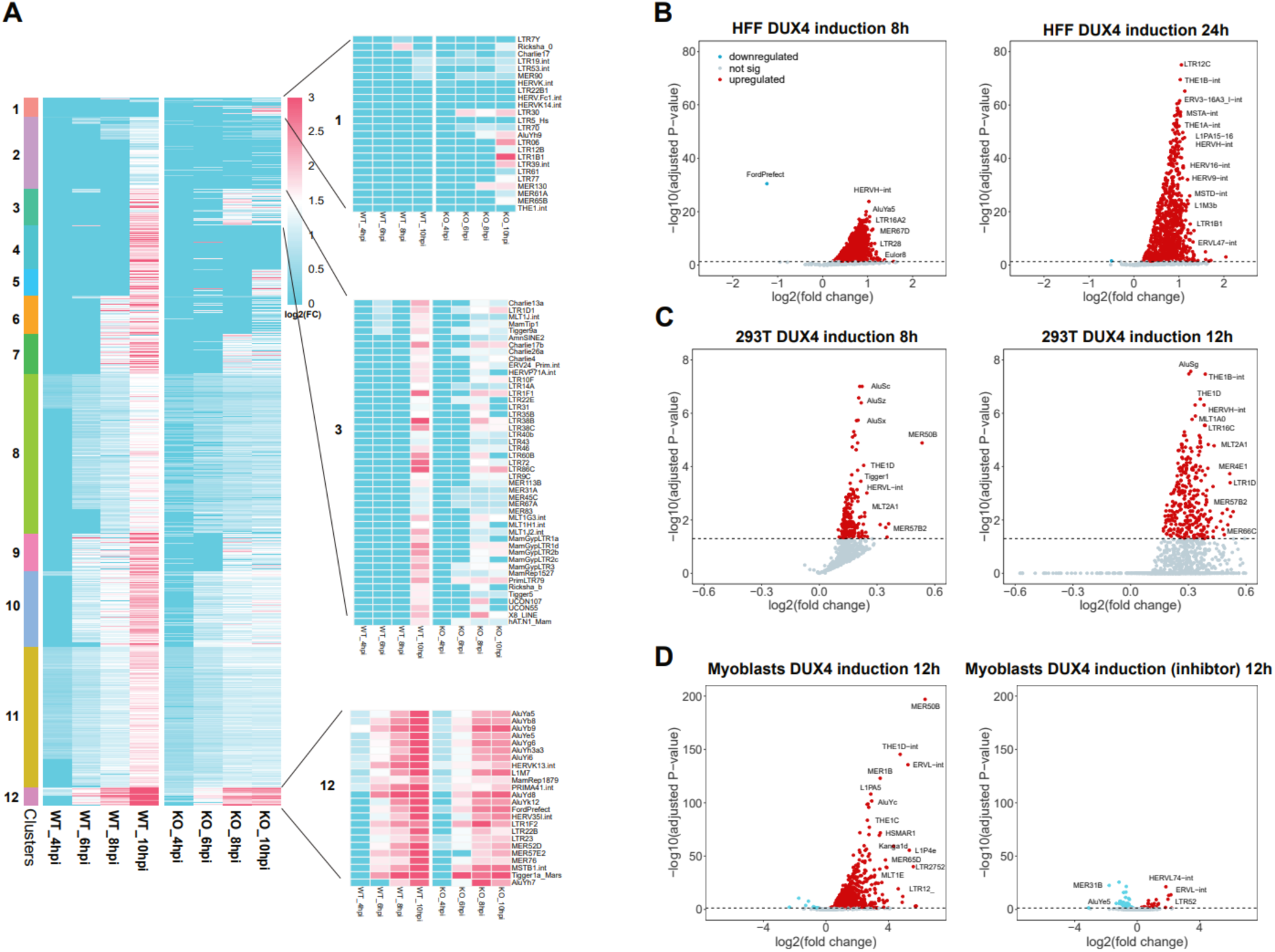
Differential TEs Regulation in Wild-Type (wt) and DUX4 Knockout (ko) Cells during HSV-1 Infection. (**A**) Heatmap depicting log2(FC) of upregulated TEs at 4, 6, 8, and 10 hours post HSV-1 infection (hpi) compared to mock in both wt and DUX4 ko HAP1 cells. TEs cluster in 12 clusters according to their expression kinetics. Values exceeding log2(FC) > 3 are capped at 3. Subclusters 1, 3, and 12 highlight TEs that are not regulated by DUX4. (**B**) Volcano plot illustrating significantly upregulated TEs upon DUX4 induction at 8 and 24 hpi in HFF cells. Red dots indicate TEs meeting the criteria of log2(FC) > 0 and padj < 0.05. (**C**) Volcano plot illustrating significantly upregulated TEs upon DUX4 induction at 8 and 12 hpi in 293T cells. Red dots indicate TEs meeting the criteria of log2(FC) > 0 and padj < 0.05. (**D**) Volcano plot illustrating significantly upregulated TEs upon DUX4 induction at 12 hours and reduction of upregulated TEs after the addition of DUX4 inhibitor in myoblasts cells. Red dots indicate TEs meeting the criteria of log2(FC) > 0 and padj < 0.05.

### KAP1 controls DUX4- and TEs-activation

KAP1 is known to be a master regulator of TEs and functions as a highly SUMOylated protein that orchestrates the assembly of the TEs silencing machinery. Silencing is mediated through its recruitment of the histone methyltransferase SETDB1 in a SUMO-dependent manner (64). Notably, prior studies have indicated that KAP1 can suppress the DUX4 homologue Dux in a LINE1-dependent fashion (65) and binding to Dux in mice (66). In addition, work by the Trono lab showed an important role of KAP1 in the regulation of HCMV latency. Our investigation shows that KAP1 specifically binds to the DUX4 locus in HFF cells upon HSV-1 infection (Fig.5A). Further analysis of CUT&RUN data revealed that another KAP1:CRAB complex member, SETDB1, as well as corresponding histone marks H3K9me3 and H3K14ac, are present at the DUX4 locus. KAP1 binding increased upon HSV-1 infection, but notably, the binding of SETDB1 and associated H3K9me3 and H3K14Ac peaks were reduced following HSV-1 infection (Fig.5A), as were those associated with TEs (Fig.5B). Intriguingly, HSV-1 infection induces KAP1 phosphorylation at serine 473 and serine 824 (Fig.5C). Mutation of serine 473 and serine 824 to alanine, in KAP1 ko cells that were reconstituted with KAP1 wt and mutants resulted in reduced TEs expression compared to wt cells upon HSV-1 infection for 8 hours (Fig.S16A, B). Phosphorylation of KAP1 at serine 824 upon HSV-1 infection is dependent on ATM activation, as it can be blocked by the ATM-inhibitor KU60019, and is therefore part of the cellular response to viral infection. In contrast, Serine 473 phosphorylation is ATM independent and directly mediated by the HSV-1 kinase US3 (Fig.5D), as shown by overexpression of US3. Serine 473 phosphorylation is known to interfere with transcriptional repression of KRAB-zinc finger protein (KRAB-ZFP) target genes, thus disrupting its function and potentially triggering the activation of DUX4 and subsequent TEs (67). To test our hypothesis, we analyzed RNA-seq datasets derived from wt and KAP1 knockdown (KD) cells (Fig.S11B, C) both before and after exposure to HCMV. Remarkably, KAP1 KD dramatically induces the expression of DUX4 target genes in the absence of infection (Fig.5E) as well as the induction of TEs (Fig.5F). Of note, the number of HCMV reads is not significantly affected by kd of KAP1 (Fig.S1C).

**Figure 5:**
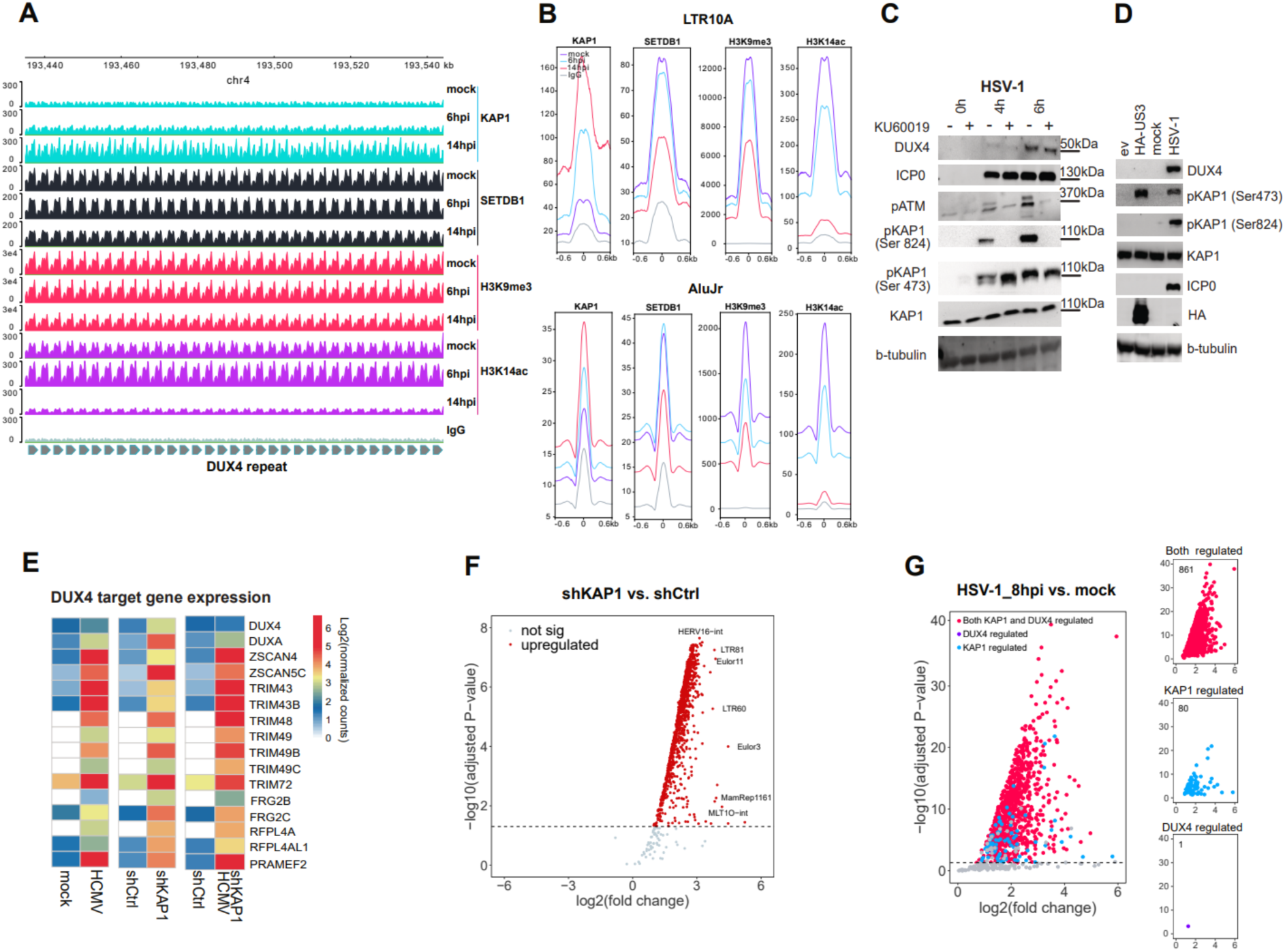
Investigating the Role of DUX4 and KAP1 in TEs activation upon herpesviral Infections. (**A**) Schematic representation and KAP1 CUT&RUN tracks showed KAP1 specifically binds to the repetitive DUX4 locus mock treated as well as HSV-1 infected cells at 6 and 14 hpi. SETDB1 and the histone marker H3K9me3 and H3K14ac CUT&RUN tracks showed the reduction of SETDB1 binds to DUX4 and corresponding histone modification upon HSV-1 infection at 6 and 14 hpi. (**B**) Peak count frequency of KAP1, SETDB1, and histone marker (H3K9me3 and H3K14ac) peaks overlapping with the TEs LTR10A and AluJr. (**C**) Western blot analysis of HFF cells infected with HSV-1 for 14h with MOI of 5. Including expression of DUX4 and phosphorylation of KAP1 and ATM. Tubulin serves as loading control. ICP0 serves as a control for viral infection. (**D**) Transfection of 293T cells with a plasmid encoding for HA-tagged HSV-1 kinase US3. 293T infected with HSV-1 served as control. Western Blot analysis of DUX4, KAP1, phospho-KAP1, ICP0 as infection control and tubulin as loading control. One representative experiment of n=3 is shown. (**E**) Heatmap displaying the number of normalized reads per gene (log2) with a focus on DUX4 and DUX4-target genes. Left to right panel: HFF cells infected with HCMV for 48h; HFF cells transfected with shCtrl and shKAP1; HFF cells transfected with shCtrl and shKAP1 and infected with HCMV for 48h. (**F**) Volcano plot depicting upregulated TEs in HFF cells transfected with shKAP1. Red dots represent significantly upregulated TEs from the TEs analysis (log2(FC)>0, padj < 0.05). (**G**) Volcano plot showing upregulated TEs upon HSV-1 infection at 8 hpi overlapped with DUX4-induction for 24 hours and KAP1 knockdown in HFF cells. Red dots represent TEs significantly upregulated by both KAP1 and DUX4, blue dots represent TEs upregulated by KAP1, and purple dots represent TEs upregulated by DUX4 (log2(FC)>0, padj < 0.05).

We recently showed that the viral genes ICP0 and ICP4 are sufficient for induction of DUX4 by HSV-1 (34), raising the question whether KAP1 is also involved in DUX4 induction in the context of viral infection. To this end, we infected KAP1-wt and -ko A549 cells with an HSV-1 virus with deletion of ICP0. The HSV-1-ΔICP0 induced expression of the DUX4 target gene TRIM43 was induced in KAP1-ko cells but not in wt cells, indicating that a loss of ICP0 can be compensated by ko of KAP1 (Fig. S16C). Further analysis of KAP1 kd cells with and without HCMV infection reveals increased expression of TEs in the presence of virus (Fig.S17A), underscoring the dependence of DNA virus-induced TEs activation on KAP1 and DUX4. Our results not only confirm the role of KAP1 in the regulation of TEs, but also establish a KAP1- and DUX4-dependent mechanism for herpesvirus-mediated TEs activation. Interestingly, when we examined the TEs upregulated during HSV-1 and HCMV infection, the majority were found to be KAP1 and DUX4-specific, with approximately 861 and 756 TEs regulated by both DUX4 and KAP1, respectively (Fig.5G;Fig.S17B). This interplay between KAP1, DUX4 and DNA viruses sheds light on the intricate regulatory network governing TEs expression.

### TEs are upregulated in DNA virus associated human cancers

We recently showed that the germline transcription factor DUX4 is expressed in the context of DNA virus infection, namely herpesviruses (HSV-1, HSV-2, HCMV, KSHV and EBV), papillomaviruses (HPV16) and polyomaviruses (MCPyV) (34). Along this line, we wanted to investigate whether herpesviral infections also induce TEs expression in vivo. By re-analyzing single-cell RNA-seq. data of tumor samples from EBV-positive nasopharyngeal carcinoma patients, we were able to show that TEs are upregulated in EBV-positive tumor cells (Fig.6A; Fig.S18; Supplementary Table). We noted that MLT- and THE-family transposons expression is significantly higher in malignant cells expressing viral transcripts compared to bystander cells, indicating that TEs expression is driven by the virus infection (Fig.6A; Fig.S18). Next, we investigated whether tumor cells of MCPyV-positive Merkel cell carcinoma patients and HPV-positive head and neck squamous cell carcinoma patients also have elevated TEs levels. Similarly to EBV-tumors, in Merkel cell carcinoma tumor patients, the expression of MLT and THE members is significantly upregulated in MCPyV-expressing cells compared to other cell types (Fig.6B; Fig.S19; Supplementary Table). Moreover, in head and neck squamous cell carcinoma patients, the expression of these TEs is significantly upregulated in HPV-enriched malignant cells compared to bystander cells (Fig.6C; Fig.S20; Supplementary Table). Taken together, these data suggest that TEs-expression is a common feature of human DNA virus infection *in vivo* in the context of different types of tumors. Moreover, it is known that transposable elements serve as transcription factor binding sites that can activate downstream genes. Therefore, we analyzed splicing events from TEs to downstream exons of genes. Using conservative filtering criteria, we found eight examples of genes that are activated upon infection and DUX4 overexpression (two examples shown in Fig.6D and Fig.S17C) due to DUX4 binding to a TE upstream of a gene.

**Figure 6:**
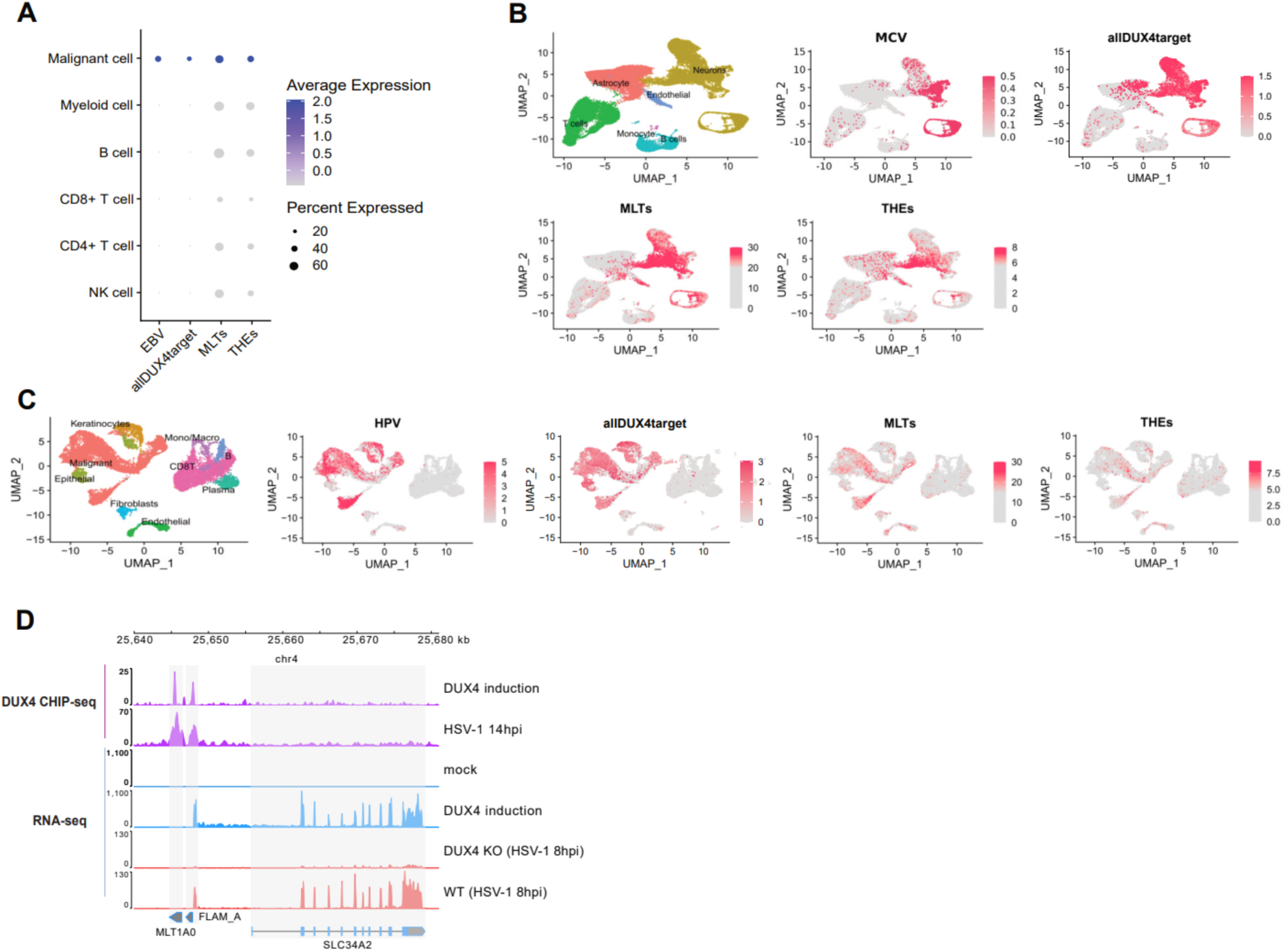
Single-Cell RNA-seq. Analysis of TEs in Virally Associated Tumors and the examples of TEs work as alternative promoters. (**A**) Single cell RNA-seq. of patient tumor tissue from Epstein-Barr Virus (EBV)-positive Nasopharyngeal Carcinoma (NPC): Dotplot showing the number of normalized EBV-specific reads per cell, normalized DUX4-target reads per cell, normalized MLTs reads per cell, and normalized THEs reads per cell in different cell types. (**B**) Single cell RNA-seq. of patient tumor tissue from Merkel Cell Polyomavirus positive Merkel Cell tumors: First panel: Cell identity map. Second panel: UMAP projection of normalized MCV-specific reads per cell. Third panel: UMAP projection of normalized DUX4-target reads per cell. Fourth panel: UMAP projection of normalized MLTs reads per cell. Fifth panel: UMAP projection of normalized THEs reads per cell. (**C**) Single cell RNA-seq. of patient tumor tissue from Head and Neck Squamous Cell Carcinoma (HNSCC) Patients positive for Human Papillomavirus (HPV): First panel: Cell identity map. Second panel: UMAP projection of normalized HPV-specific reads per cell. Third panel: UMAP projection of normalized DUX4-target reads per cell. Fourth panel: UMAP projection of normalized MLTs reads per cell. Fifth panel: UMAP projection of normalized THEs reads per cell. (**D**) Schematic representation and CHIP-seq. tracks illustrate the binding of DUX4 to the TEs MLT1A0 and FLAM_A that are located in the promoter region of the gene SLC34A2 after DUX4 induction and HSV-1 infection as well as RNA-seq. tracks of the new transcript of gene SLC34A2 after DUX4 induction and HSV-1 infection.

## Discussion

The relationship between hosts and viruses involves ongoing co-evolution, with viruses exerting persistent selection pressure that drives the evolution of antiviral mechanisms within host cells. Our comprehensive transcriptomic analysis of human cells infected with HSV-1 and HCMV reveals a noteworthy upregulation of TEs post infection. This observation is further validated through single-cell RNA-seq. analysis of patients with MCPyV, HPV, and EBV-associated cancers *in vivo*, emphasizing the clinical relevance of these findings. Despite previous reports indicating that viral infections can impact TEs expression (64, 68), the exact molecular mechanisms underlying this widespread impact remained elusive until now. A pivotal discovery of our study is the identification of the germline pioneer transcription factor DUX4 as a key player in the activation of TEs induced by DNA viruses. Utilizing DUX4 ChIP-seq., we found a significant association between virus-induced expression of DUX4 and expression of TEs, with 69% of DUX4 binding sites occurring at TEs. The functional relevance of DUX4 is underscored by experiments demonstrating that its overexpression significantly induces TEs expression, while its knockout diminishes TEs expression upon HSV-1 infection. In addition, we found a strong association of DUX4 binding with opening of chromatin at DUX4 binding sites, indicated by ATAC-Seq. peaks and reduction of H3K9-trimethylation. This positions DUX4 as a central regulator in the expression of TEs during DNA viral infections.

In prior research, an important role of SUMOylation-modified KAP1 in TEs-silencing has been demonstrated, triggering the establishment of repressive histone marks at TEs sites. Notably, the SUMOylated form of KAP1 is rapidly lost following Influenza A virus (IAV) infection (71), most likely due to the phosphorylation of KAP1 (72). Phosphorylation of KAP1 is known to regulate binding to heterochromatin protein 1 (HP1), KRAB-ZFP and the histone methyltransferase SETDB1. Our investigation extends these findings by demonstrating the phosphorylation of KAP1 during HSV-1 infection at two different residues, one mediated by ATM-signaling and the second one mediated by the viral kinase US3. ChIP-seq. as well as CUT&RUN analysis shows binding of KAP1 to the DUX4 locus and KAP1-ko induced DUX4 target gene- and TEs-expression, suggesting a regulatory role of KAP1 in modulating DUX4 and subsequent expression of TEs. This goes along with reduced CUT&RUN signals for SETDB1, H3K9me3 and H3K14Ac upon HSV-1 infection, both at the DUX4 locus and at loci for activated TEs. Our discovery demonstrates a novel pathway by which DNA viruses, exemplified by HSV-1, may induce the activation of TEs through the phosphorylation of KAP1 and subsequent modulation of DUX4.

The interplay between viruses and their hosts includes the reactivation and co-optation of TEs for effective antiviral defense. In general, it is thought that the activation of TEs results in the activation of innate immune pathways because TEs are derived from viruses and therefore still contain pathogen associated molecular patterns (PAMPs) that might be recognized by pattern recognition receptors (PRRs). Activation of PRRs results in triggering cellular defense mechanisms e.g. expression of interferon stimulated genes and production of interferons that act on neighboring cells and activate antiviral defense mechanisms. However, we showed that activation of DUX4 is directly mediated by herpesviral proteins; for HSV-1 two immediate-early proteins, ICP0 and ICP4 are sufficient to induce expression of DUX4 (34). In addition, the HSV-1 kinase US3 phosphorylates KAP1 at Ser473, thereby lifts restriction by KAP1, which results in TEs expression. This would argue against an antiviral mechanism since viruses generally avoid activation of antiviral mechanisms by all means. In addition, we previously showed that DUX4 activation is critical for efficient replication of HSV-1 and HSV-2 (34). Therefore, it might be conceivable that activation of TEs has beneficial effects for the replication or persistence of DNA viruses. Members of herpesviruses, papillomaviruses and polyomaviruses are known to establish persistent infections that lead to oncogenic transformation of cells in a proportion of cases, and we could demonstrate that TEs are activated in cancer cells containing viral transcripts from patients (Fig.6A, B, C). In this context, our present investigation significantly contributes to the understanding of the complex relationship between DNA viruses and the host genome in respect to TEs-activation.

DNA virus genomes contain multiple genes that are derived from host genes. For herpesviruses, some of the genes are conserved across all human herpesviruses, like the large ribonucleotide reductase subunit, whereas others are only present in individual subfamilies like the viral FGARATs of gamma-herpesviruses KSHV and EBV (73). It is thought that the genes were “stolen” by viruses at one point during evolution by horizontal gene transfer and then diversified during the coevolution with the respective hosts. The viral FGARATs for example show about 25% sequence identity with human FGARAT and have lost their role in nucleotide metabolism (74). However, viral FGARATs have evolved as potent inhibitors of the antiviral function of PML nuclear bodies (75, 76). Host gene homologues are also present in the genome of Poxviruses. Poxviruses are also DNA viruses, but in contrast to herpesviruses, they replicate in the cytoplasm. Two recent papers showed that the uptake of host genes into the poxvirus genome is mediated by retrotransposons (77, 78). The authors showed that mRNA can be reverse transcribed by the reverse transcriptase of LINE-1 elements and then integrate into poxvirus genomes. Although the exact mechanism might be different between nuclear and cytoplasmic replicating viruses due to different replication strategies, it shows an important role for retrotransposons in the host adaptation of DNA viruses. It is conceivable that the induction of retrotransposons is crucial for the evolution of DNA viruses and subsequent studies will be needed to investigate which role TEs activation plays in DNA virus infection and viral oncogenesis in detail.

The results presented here not only provide a characterization of TEs activation during DNA virus infection but also highlight the essential roles of DUX4 and KAP1 as key regulators in this process. The intricate molecular mechanisms uncovered in this study offer valuable insights into the dynamics of host genome-virus interactions. Understanding the role and regulation of TEs during viral infections is crucial, as TEs are integral components of cis-regulatory elements, and their dysregulation can have profound implications for host genomic stability and function. These findings have implications for broader fields, including virology and cancer biology, as the study includes analysis of viral-associated cancers. The identification of DUX4 as a key mediator in TEs activation suggests potential therapeutic targets for controlling TEs expression during viral infections, with potential implications for diseases associated with viral etiology. Overall, this study significantly advances our knowledge of the molecular mechanisms governing TEs activation in the context of DNA virus infections, opening avenues for further research and therapeutic exploration.

## Data Availability

The raw sequencing datasets have been uploaded to the Gene Expression Omnibus (GEO) database, including RNA-Seq of DUX4 over-expression in HFF and 293T cells, HCMV (BAC2) and KSHV infection, CUT&RUN of KAP1, SETDB1, H3K9me3 and H3K14ac (GSE280380), RNA-Seq of HCMV (TB40/E) infection and KAP1 knockdown (GSE256111), and RNA-seq of DUX4-ko (GSE287622). The study used publicly available datasets from GEO, including bulk RNA-seq of HSV-1 infection (GSE59717) (61), KSHV infection (GSE123898) (79), EBV infection (GSE155345) (80), and DUX4 induction and inhibition in myoblasts cells (GSE130522) (63). ATAC-seq of HSV-1 infection (GSE100611) (81), CHIP-seq of H3K9me3 and H3K27me3 upon HSV-1 infection (GSE124803) (62), CHIP-seq of DUX4 (GSE33838 and GSE174759) (29, 34), as well as the single-cell RNA-seq data of nasopharyngeal carcinoma (GSE162025) (82), Head and neck squamous cell carcinoma (GSE164690) (83) and Merkel cell carcinoma tumor (GSE226438) (84). UCSC genome browser session links for the ATAC-seq. and ChIP-seq. data are available under the following link: https://genome-euro.ucsc.edu/s/tanjiang/TEs_track.

## Acknowledgements

We are grateful to Dr. Daniel Sauter (Tübingen, Germany), Dr. Andrea Thoma-Kress (Erlangen, Germany) and Prof. Ben Hale (Zurich, Switzerland) for providing plasmids. This study was supported by the BMBF through Junior Research Group “Duxdrugs” (BMBF 01KI2017, F.F.) and the Deutsche Forschungsgemeinschaft through project DFG En 423/5-1 (to A.E.), in the framework of Research Unit FOR5200 DEEP-DV (443644894) through project STA357/8-1 to T.S. and project LA 2941/18-1 to M.L.. E.N. is a member of the Spemann graduate school of biology and medicine (SGBM).

## Declaration of interests

The authors declare no competing interests.

---

**Fig. S1.**
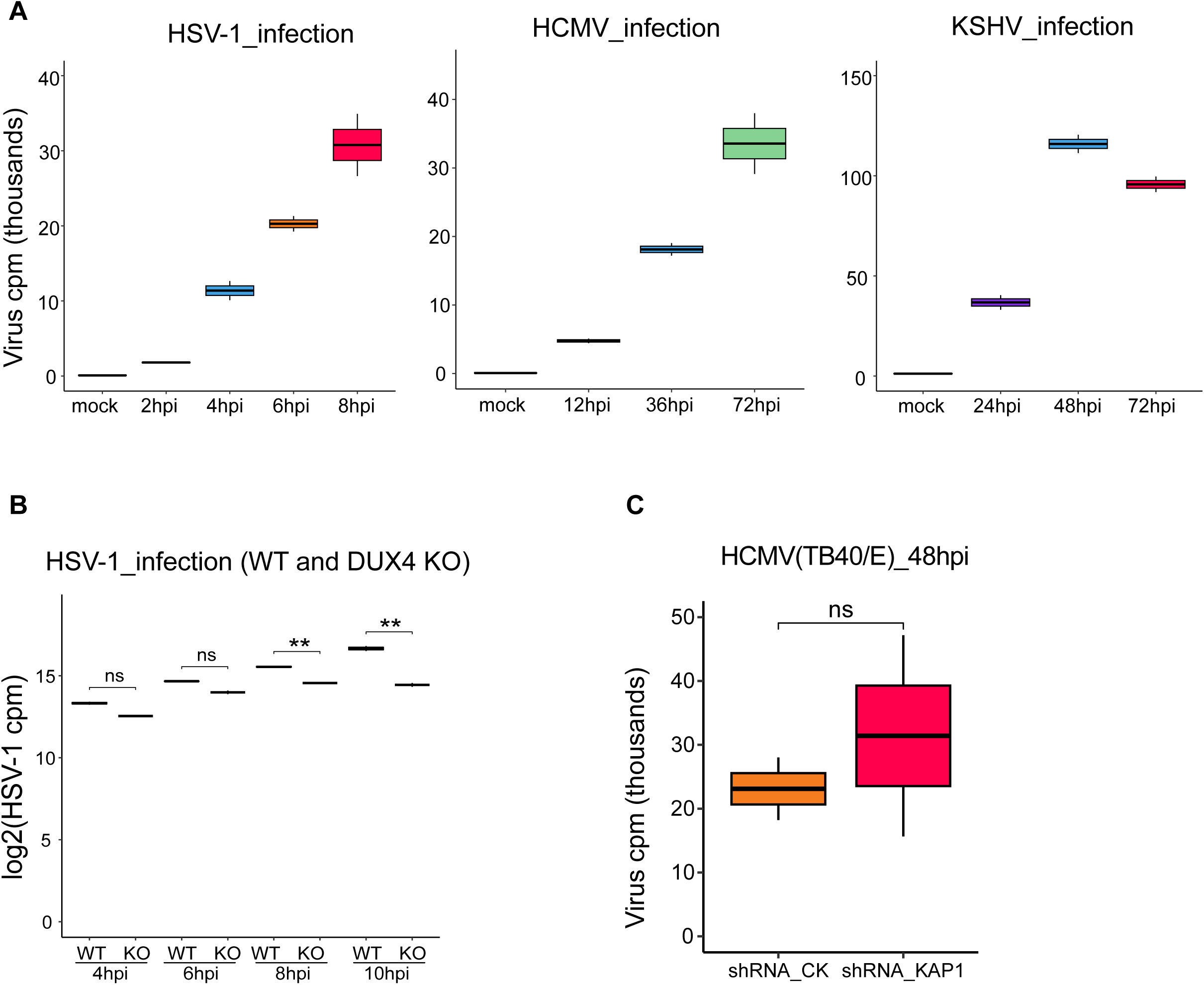
(**A**) Box plots showing the normlized virus counts per million (cpm) in mock cells and upon HSV-1 infection at 2, 4, 6, and 8 hours (hpi), HCMV infection at 12, 36 and 72 hours (hpi) and KSHV infection at 24, 48 and 72 hours (hpi). (**B**) Box plot showing the log2 normlized virus counts per million (cpm) upon HSV-1 infection at 4, 6, 8 and 10hpi in WT and DUX4 KO of HAP1 cells. (**C**) Box plot showing the normlized virus counts per million (cpm) upon HCMV(TB40/E) infection at 48hpi in control and KAP1 knock down cells. Statistical significance was determined using two-sided nonparametric Wilcoxon rank-sum tests (ns: not significant, *P < 0.05, **P < 0.01, ***P < 0.001).

**Fig. S2.**
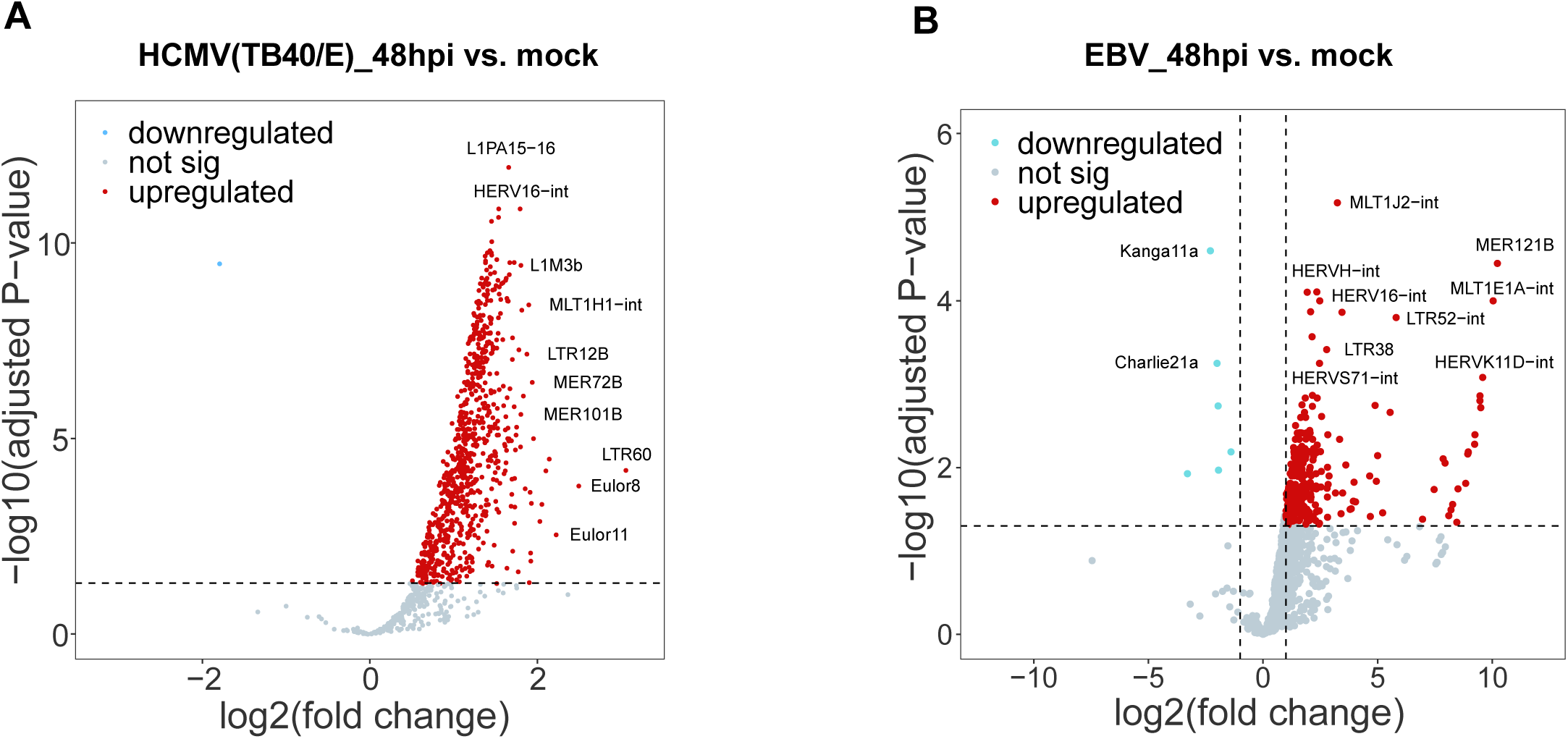
(**A**) Volcano plot showing significantly upregulated TEs upon HCMV strain TB40/E infection at 48 hpi in HFF cells. Red dots indicate TEs meeting the criteria of log2(FC) > 0 and padj< 0.05. (**B**) Volcano plot showing significantly changed TEs of upon EBV infection at 48 hpi in B cells. Red dots indicate TEs meeting the criteria of log2(FC) > 1 and padj< 0.05 and blue dots indicate TEs meeting the criteria of log2(FC) < -1 and padj< 0.05.

**Fig. S3.**
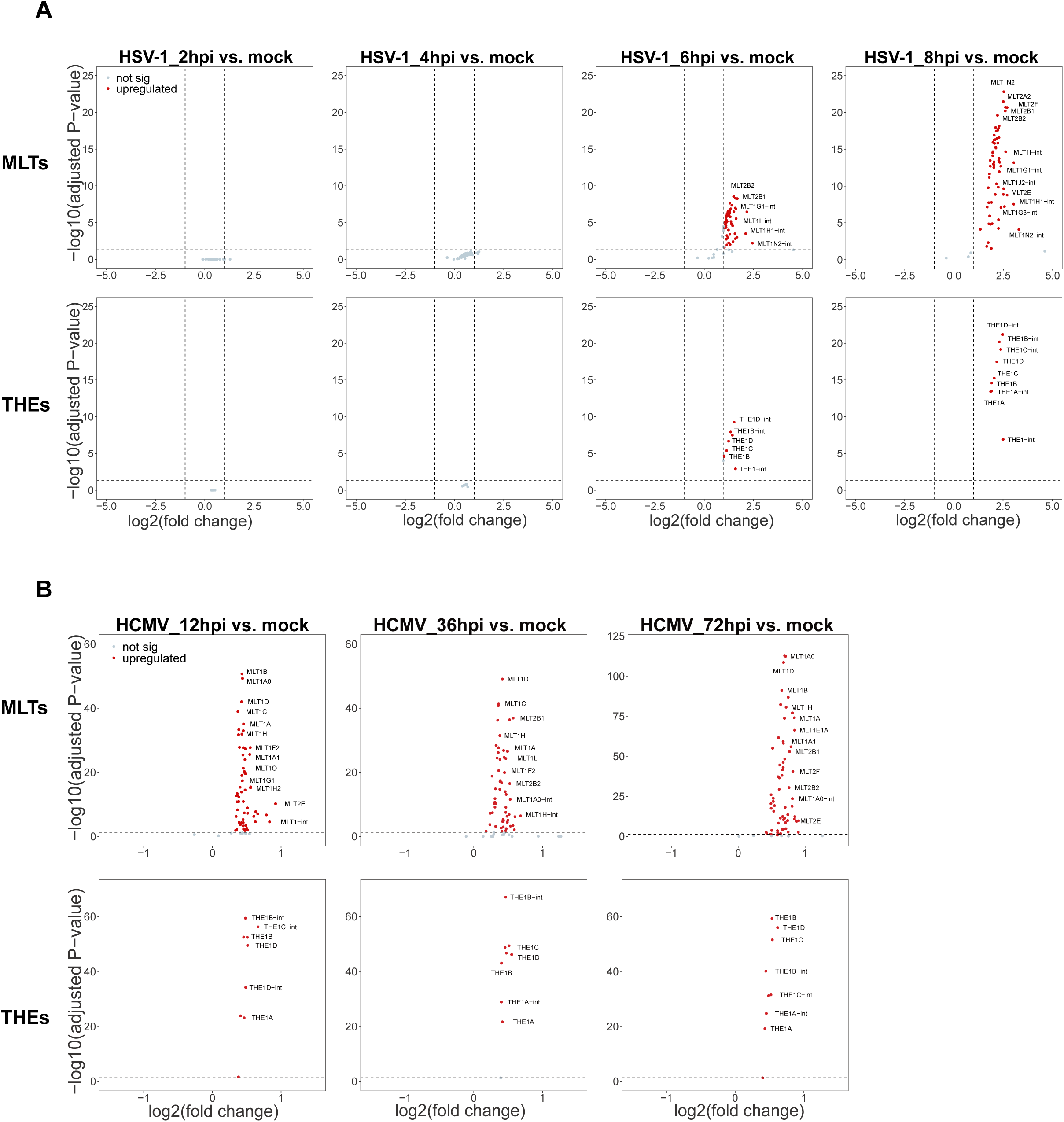
(**A**) Volcano plot showing significantly upregulated TEs of the MLT- and THE-families upon HSV-1 infection at 2, 4, 6 and 8 hpi in HFF cells. Red dots indicate TEs meeting the criteria of log2(FC) > 1 and padj < 0.05. (**B**) Volcano plot showing significantly upregulated MLTs and THEs upon HCMV infection at 12, 36 and 72hpi in HFF cells. Red dots indicate TEs meeting the criteria of log2(FC) > 0 and padj < 0.05.

**Fig. S4.**
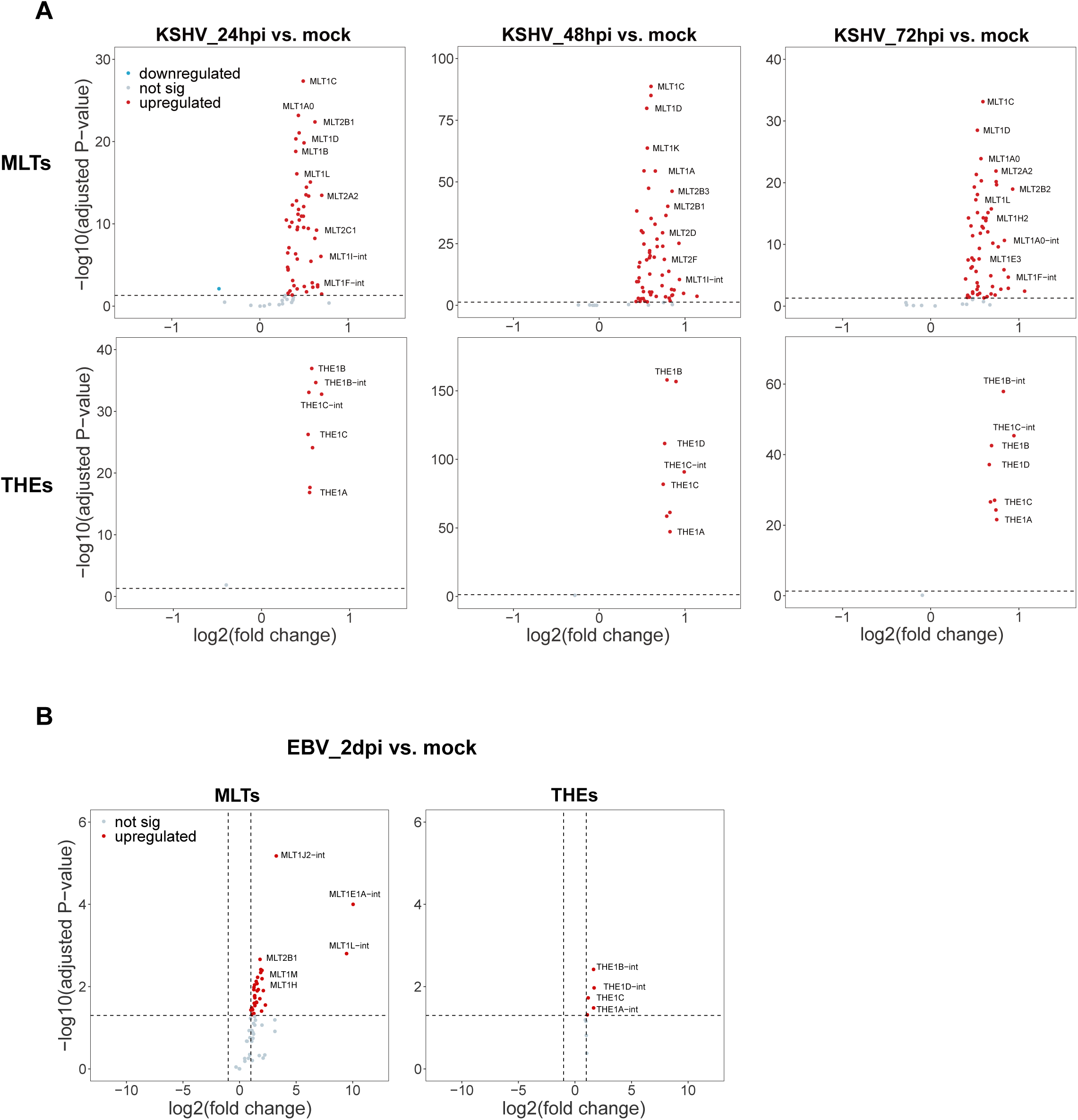
(**A**) Volcano plot showing significantly upregulated TEs of the MLT- and THE-families upon KSHV infection at 24, 48 and 72hpi in HFF cells. Red dots indicate TEs meeting the criteria of log2(FC) > 0 and padj < 0.05. (**B**) Volcano plot showing significantly upregulated MLTs and THEs upon EBV infection at 2 dpi in B cells. Red dots indicate TEs meeting the criteria of log2(FC) > 1 and padj < 0.05.

**Fig. S5.**
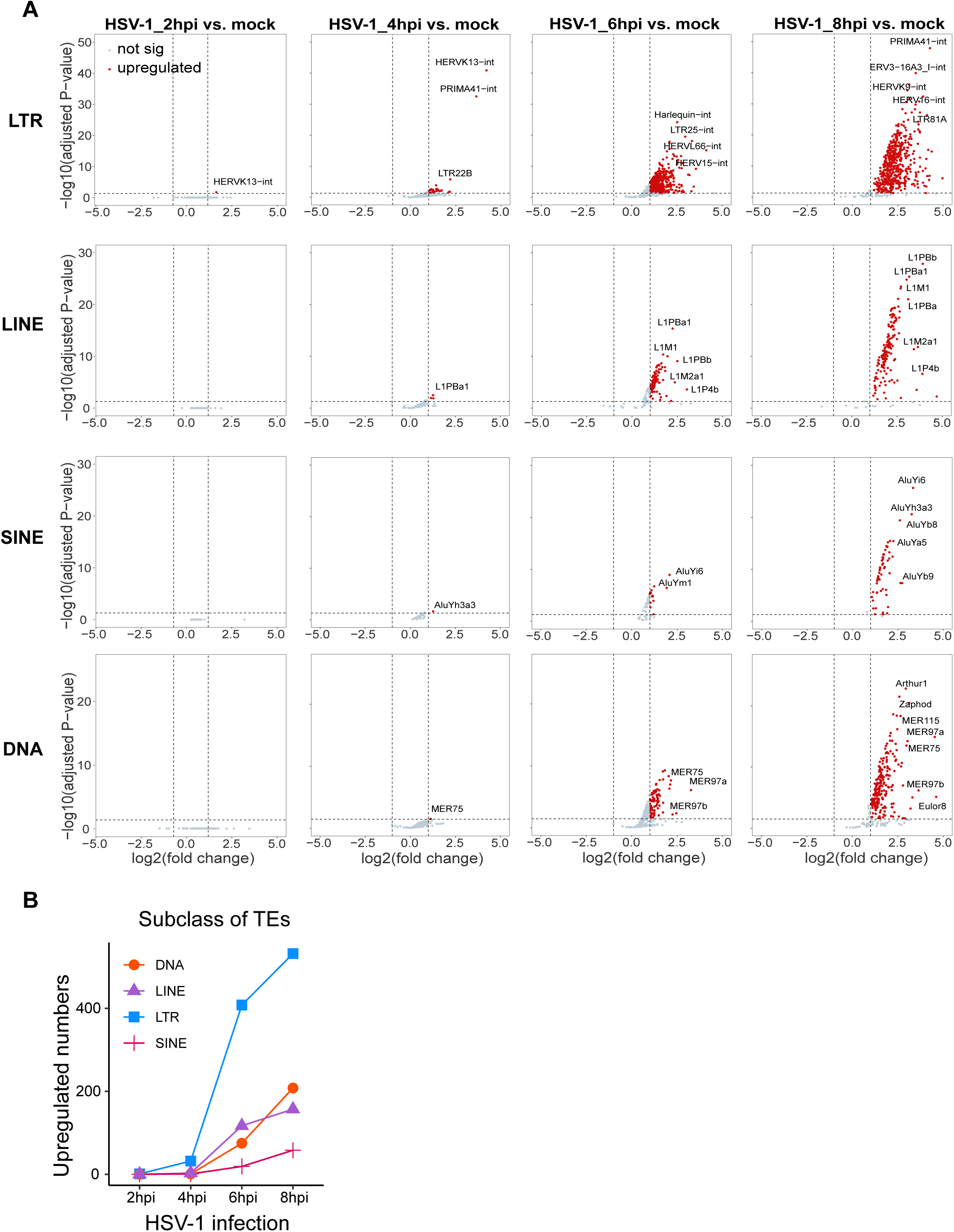
(**A**) Volcano plot showing significantly upregulated TEs of 4 subclasses (LTR, LINE, SINE and DNA respectively) upon HSV-1 infection at 2, 4, 6, and 8 hpi in HFF cells. Red dots indicate TEs meeting the criteria of log2(FC) > 1 and padj< 0.05. (**B**) Line plot showing the number of upregulated TEs of 4 subclasses.

**Fig. S6.**
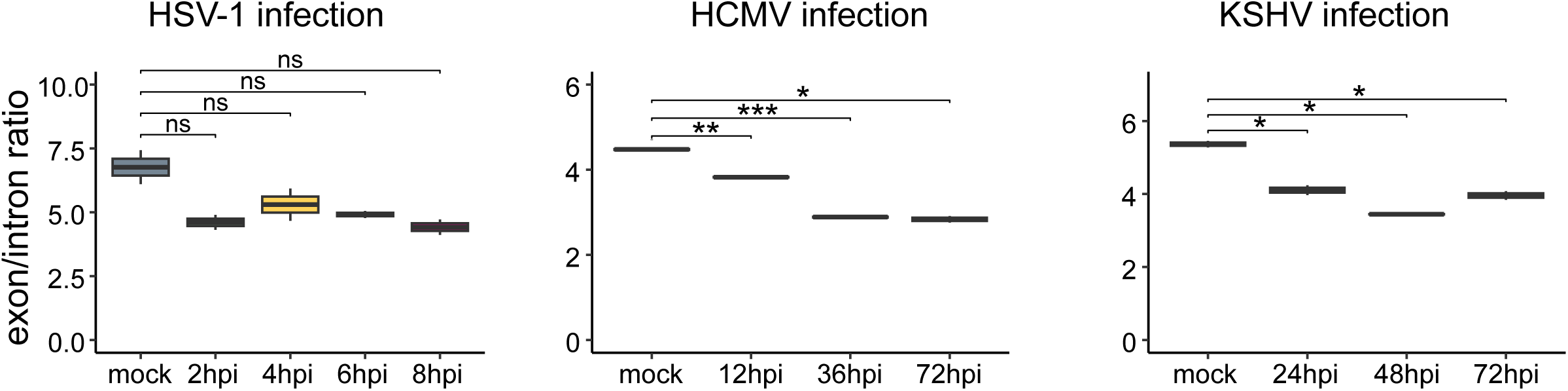
Box plots showing the ratio of reads mapping to exons and introns in mock cells and upon HSV-1 infection at 2, 4, 6, and 8 hours (hpi), HCMV infection at 12, 36 and 72 hours (hpi) and KSHV infection at 24, 48 and 72 hours (hpi). Statistical significance was determined using two-sided nonparametric Wilcoxon rank-sum tests (ns: not significant, *P < 0.05, **P < 0.01, ***P < 0.001).

**Fig. S7.**
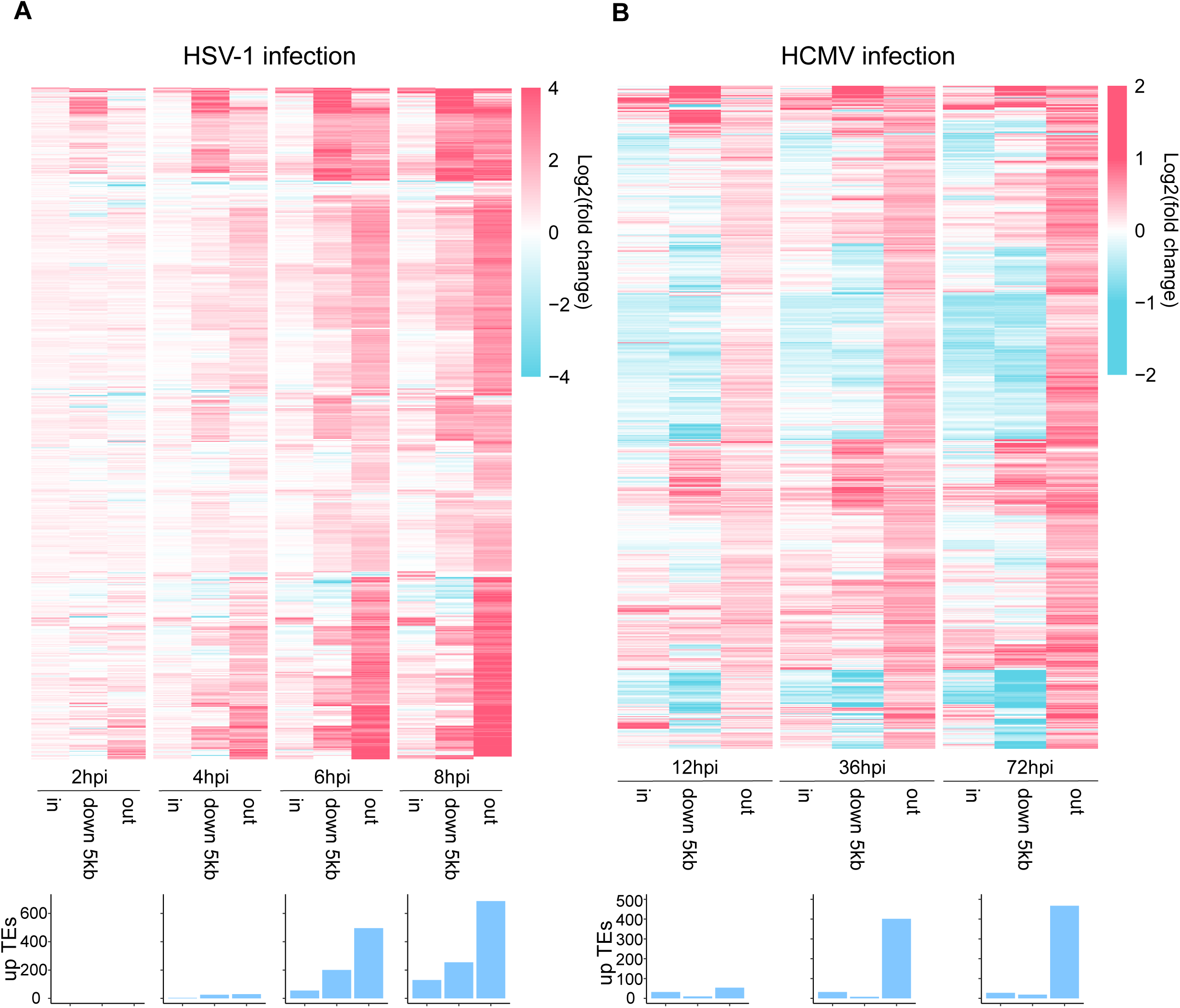
(**A**) Heatmap depicting the log2(FC) of upregulated TEs in whole genome analysis that are located within genes (in), 5 kb downstream of genes (down 5kb) and more than 5kb outside of genes (out) at 2, 4, 6, and 8 hours post HSV-1 infection (hpi) compared to mock. Barplots show the number of upregulated TEs meeting the criteria of log2(FC) > 0 and padj < 0.05 in corresponding location at corresponding time points. Values exceeding log2(FC) > 4 are capped at 4. (**B**) Heatmap depicting the log2(FC) of upregulated TEs in whole genome analysis that are located within genes (in), 5 kb downstream of genes (down 5kb) and more than 5kb outside of genes (out) at 12, 36 and 72 hours post HCMV infection (hpi) compared to mock. Barplot shows the number of upregulated TEs meeting the criteria of log2(FC) > 0 and padj < 0.05 in corresponding location at corresponding time points. Values exceeding log2(FC) > 2 are capped at 2.

**Fig. S8.**
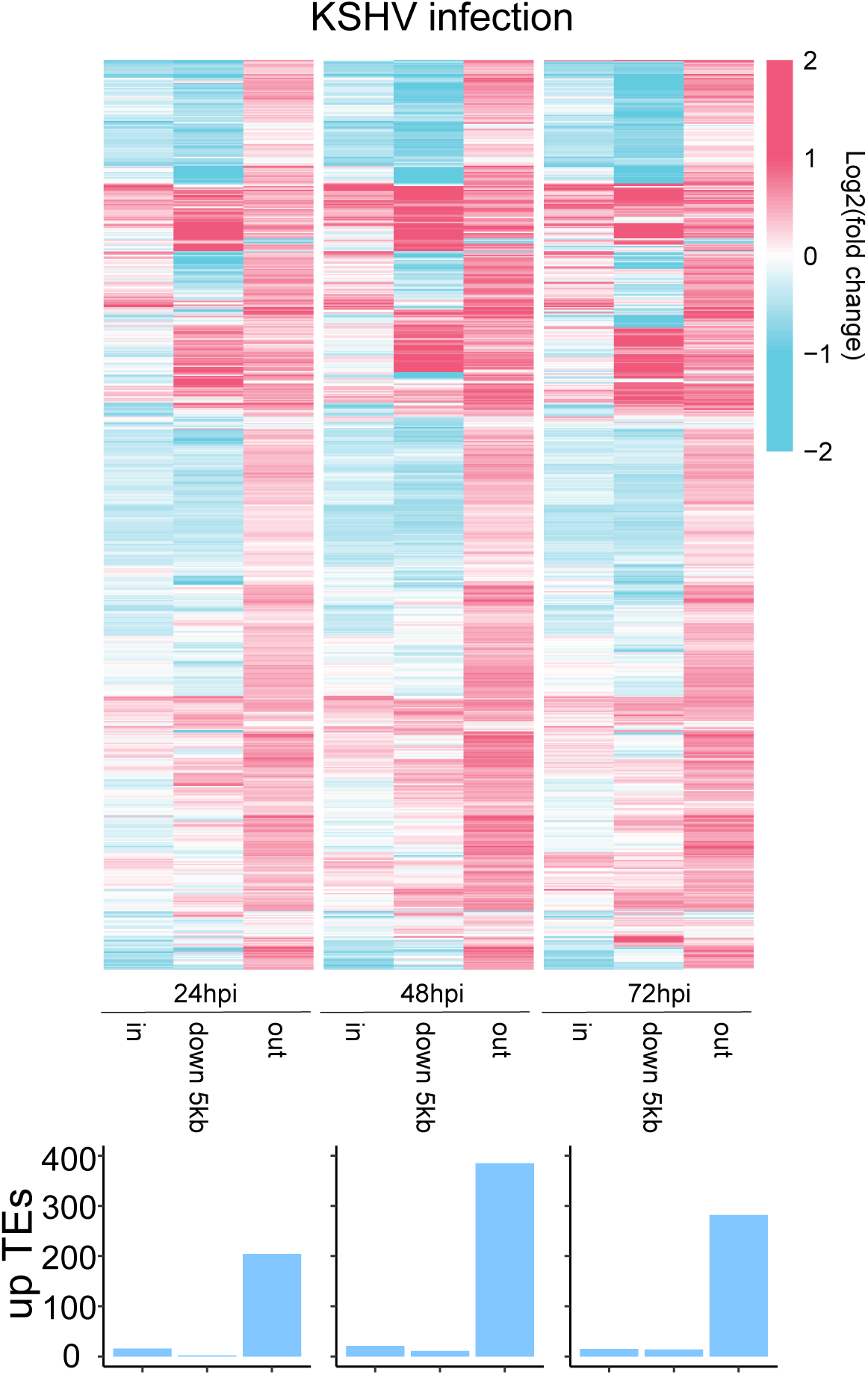
Heatmap depicting the log2(FC) of upregulated TEs in whole genome analysis that are located within genes (in), 5 kb downstream of genes (down 5kb) and outside of genes (out) at 24, 48 and 72 hours post KSHV infection (hpi) compared to mock. Barplots show the number of upregulated TEs meeting the criteria of log2(FC) > 0 and padj < 0.05 in corresponding location at corresponding time points. Values exceeding log2(FC) > 2 are capped at 2.

**Fig. S9.**
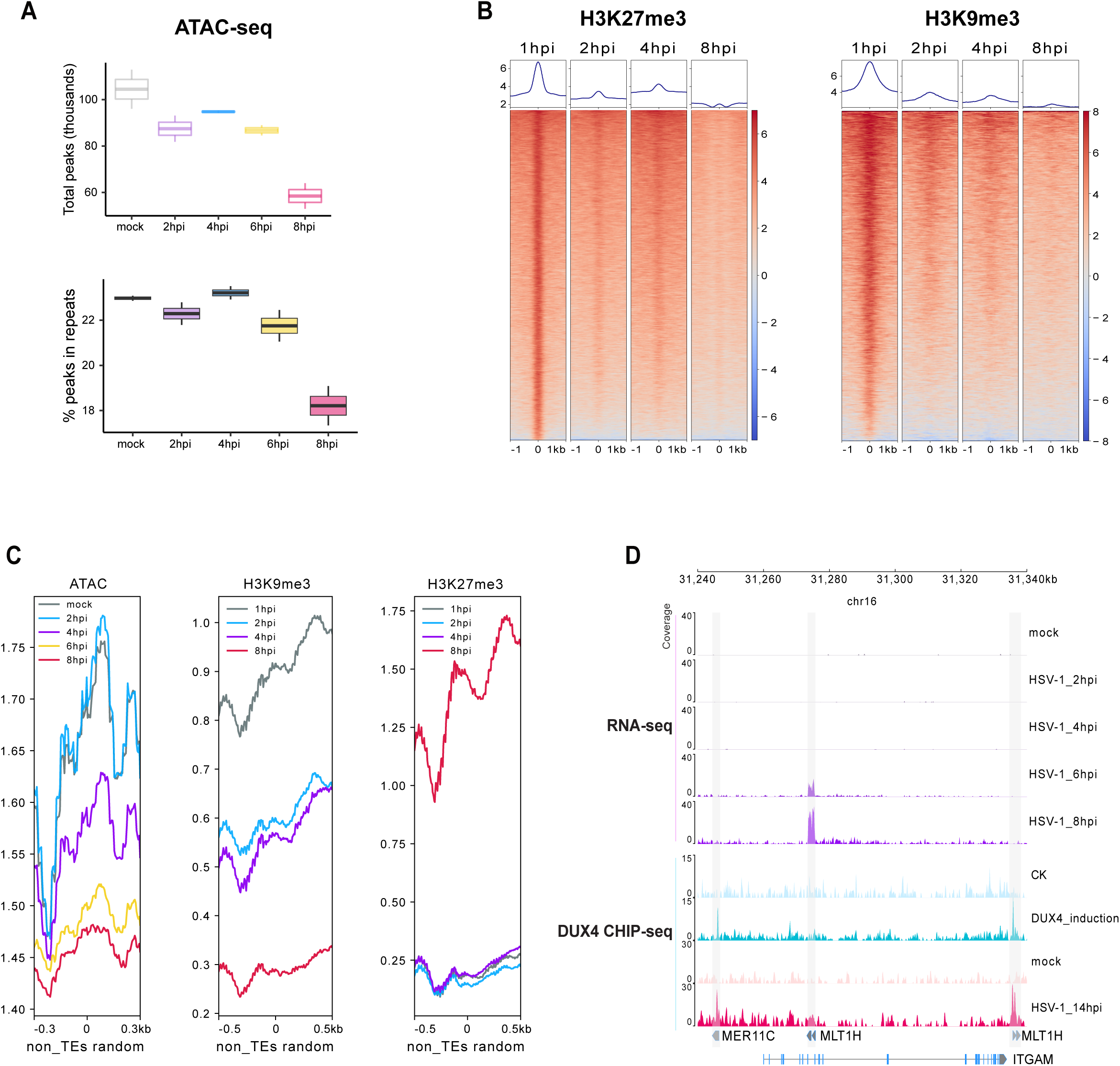
(**A**) Number of peak regions detected in ATAC-seq. of both mock and HSV-1-infected HFF cells (upper panel) and proportion of ATAC-seq peaks that overlap with repeat regions (lower panel). (**B**) Heatmap displaying enrichment levels of TEs with downregulated H3K9me3 and H3K27me3 markers upon HSV-1 infection at 1, 2, 4 and 8 hours post-infection (hpi) in HFF cells. **(C**) Peak count frequency of ATAC-seq as well as ChIP-seq peaks of transcriptional repression marks (H3K9me3 and H3K27me3) overlapping with 100000 random non-TE regions. (**D**) Schematic representation and RNA-seq tracks illustrate the expression dynamics of the TEs MLT1H and the gene ITGAM during HSV-1 infection at mock, 2, 4, 6, and 8 hpi. Schematic representation and CHIP-seq tracks illustrate the binding of DUX4 to the TEs MER11C and MLT1H after DUX4 induction and HSV-1 infection.

**Fig. S10.**
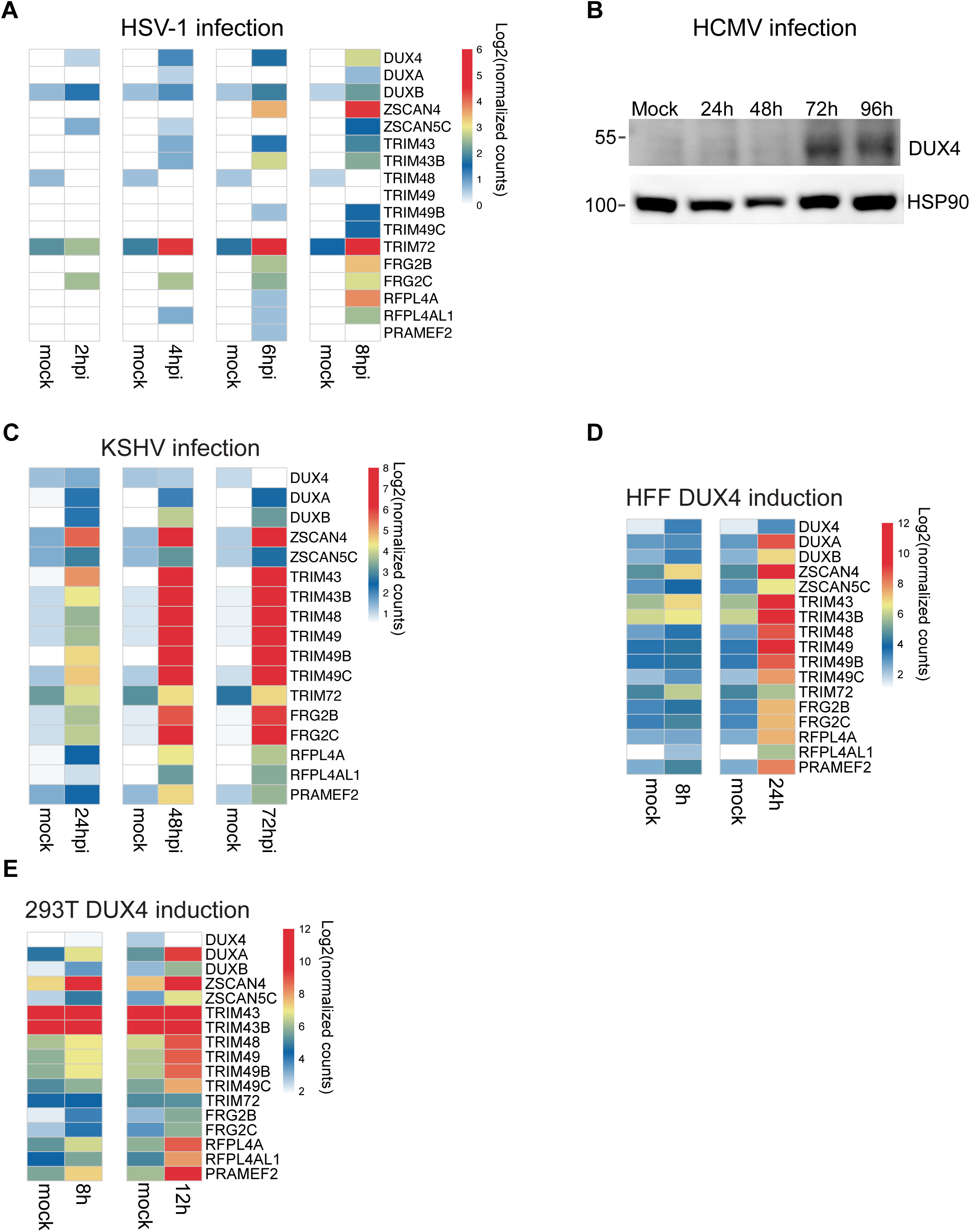
Heatmaps displaying the number of normalized counts per gene (log2) with a focus on DUX4 and DUX4-target genes or Western blot analysis of DUX4. (**A**) HSV-1 infection at 2, 4, 6, and 8 hours (hpi), (**B**) Western blot analysis of DUX4 upon HCMV infection at 24, 48, 72 and 96 hours (hpi), (**C**) KSHV infection at 24, 48 and 72 hours (hpi), (**D**) DUX4 induction for 8 and 24 hours in HFF cells, (**E**) DUX4 induction for 8 and 12 hours in 293T cells.

**Fig. S11.**
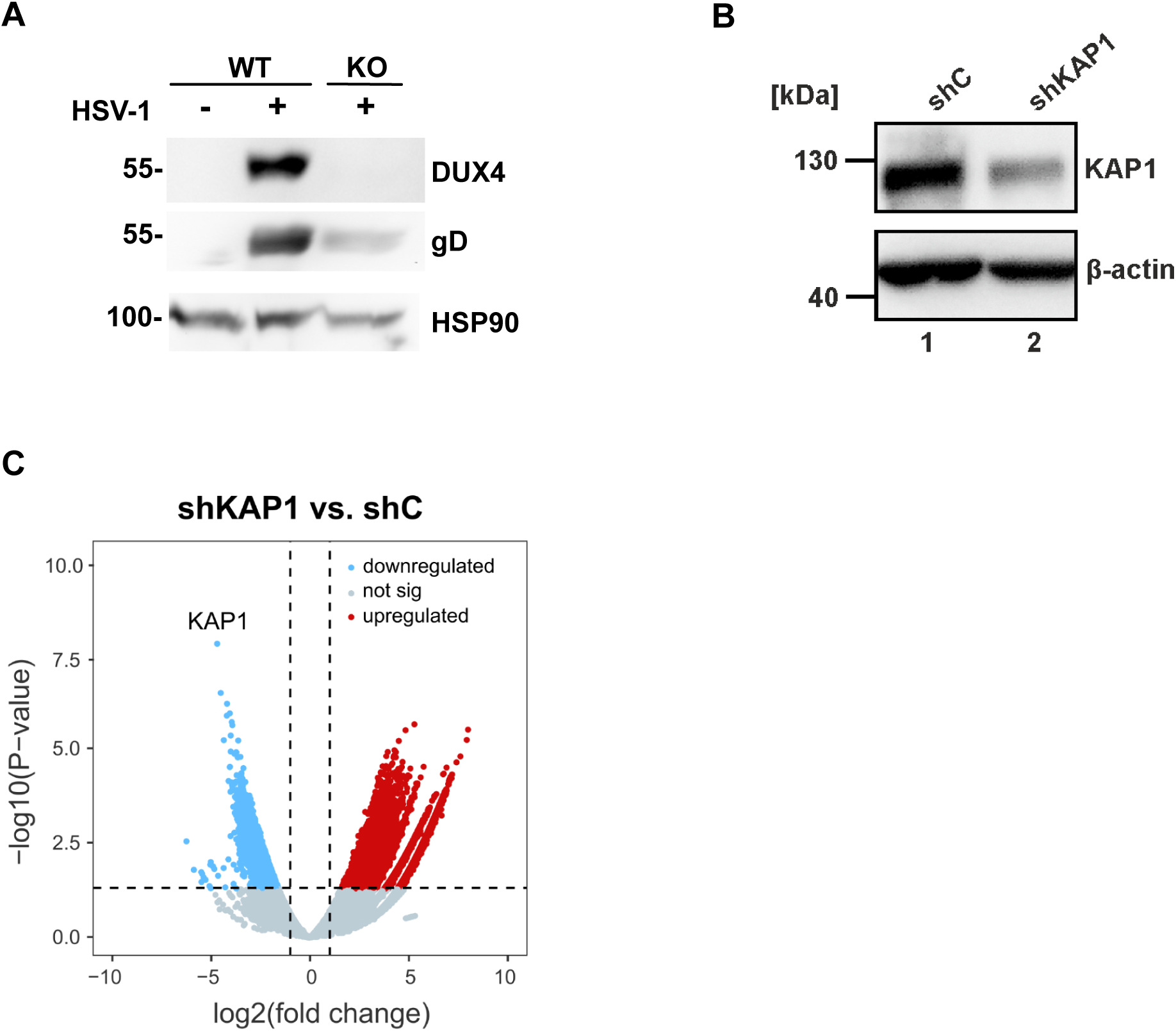
(**A**) Western blot analysis of DUX4 knockout cells (KO) and wildtype (WT) HAP1 cells. Cells were infected with HSV-1-GFP for 20h. HSP90 actin serves as loading control. **(B)** Western blot analysis of KAP1 knockdown (shKAP1) and control (shC). β actin serves as loading control. **(C)** Volcano plot showing significantly downregulated KAP1 upon KAP1 KD compared to control. Blue dots indicate the expression meeting the criteria of log2(FC) < -1 and padj < 0.05. Red dots indicate the expression meeting the criteria of log2(FC) > 1 and padj < 0.05.

**Fig. S12.**
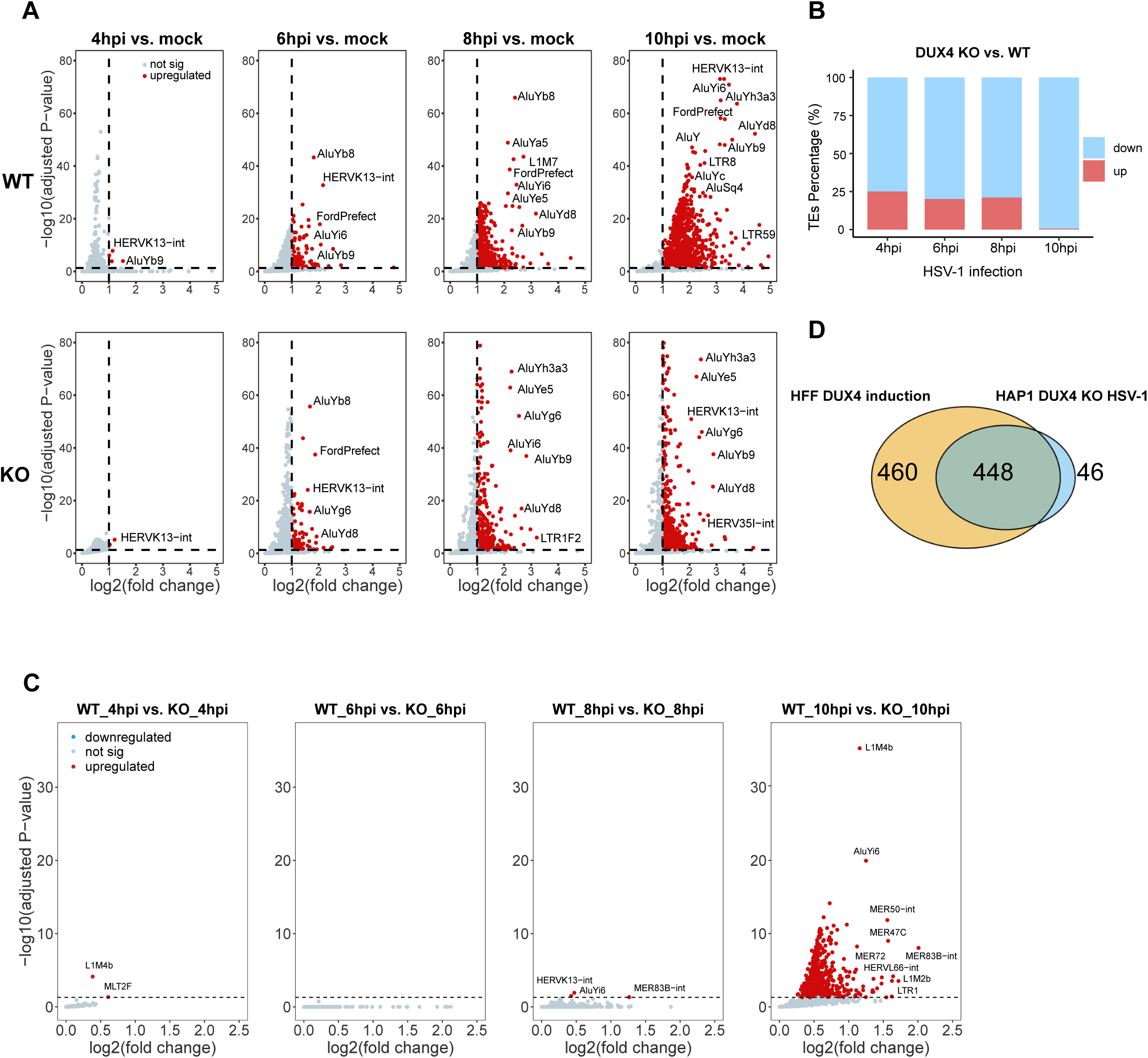
(**A**) Volcano plot illustrating significantly upregulated TEs upon HSV-1 infection at 4, 6, 8, and 10 hpi in WT and DUX4 KO. Red dots indicate TEs meeting the criteria of log2(FC) > 1 and padj < 0.05. (**B**) Percentage of up and downregulated TEs in DUX4-KO compared to DUX4-WT cells during HSV-1 infection at 4, 6, 8, and 10 hpi. (**C**) Volcano plot illustrating significantly upregulated TEs in WT compared to DUX4 KO upon HSV-1 infection at 4, 6, 8, and 10 hpi. Red dots indicate TEs meeting the criteria of log2(FC) > 0 and padj < 0.05. (**D**) Venn diagram illustrating the overlap of upregulated TEs in HFF upon DUX4 induction for 24h and downregulated TEs in HAP1 DUX4 knockout upon HSV-1 infection for 10h.

**Fig. S13.**
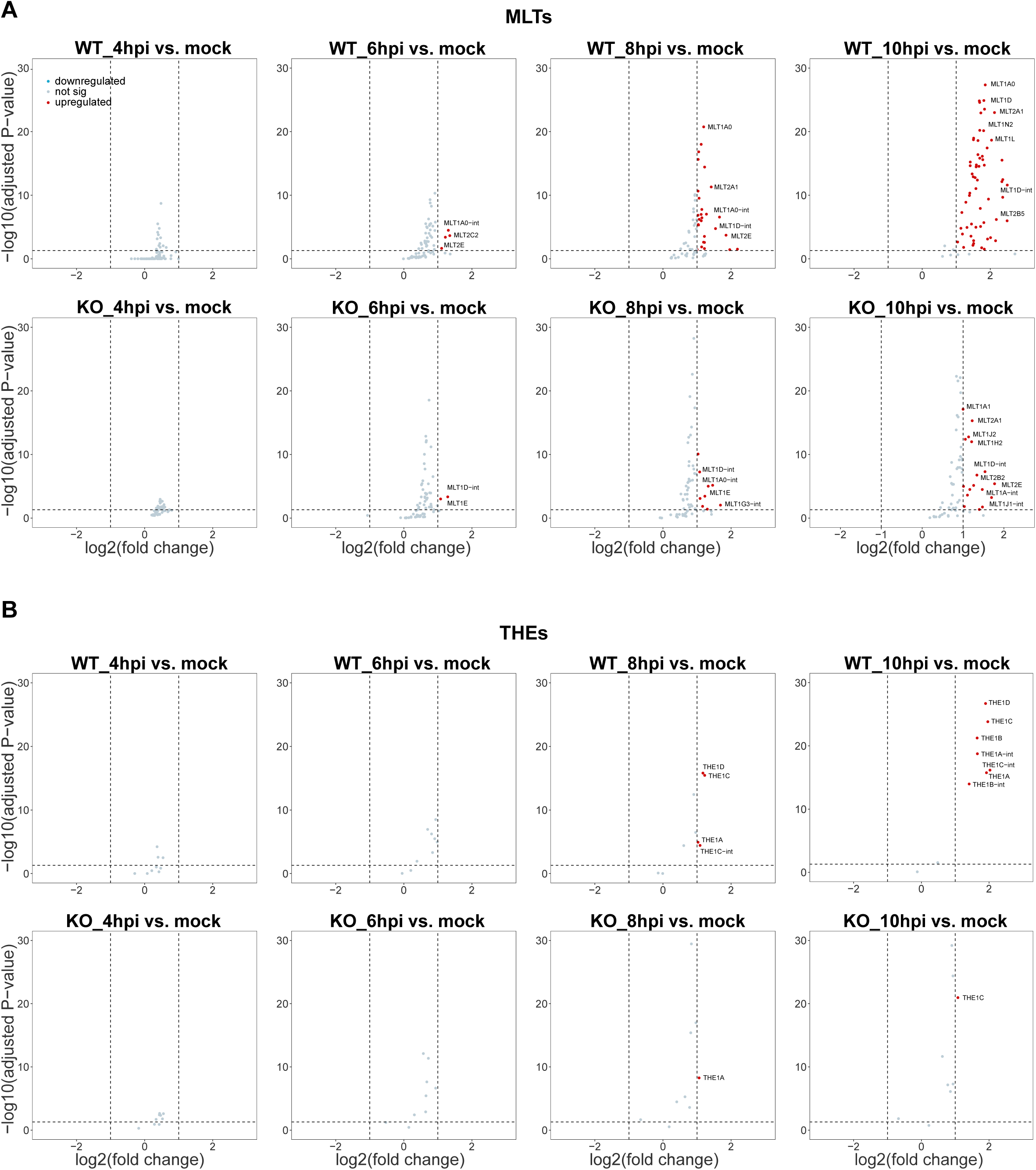
Volcano plot showing significantly upregulated TEs of the MLT- (**A)** and THE- (**B**) families upon HSV-1 infection at 4, 6, 8 and 10hpi in WT and DUX4 KO of HAP1 cells. Red dots indicate TEs meeting the criteria of log2(FC) > 1 and padj < 0.05.

**Fig. S14.**
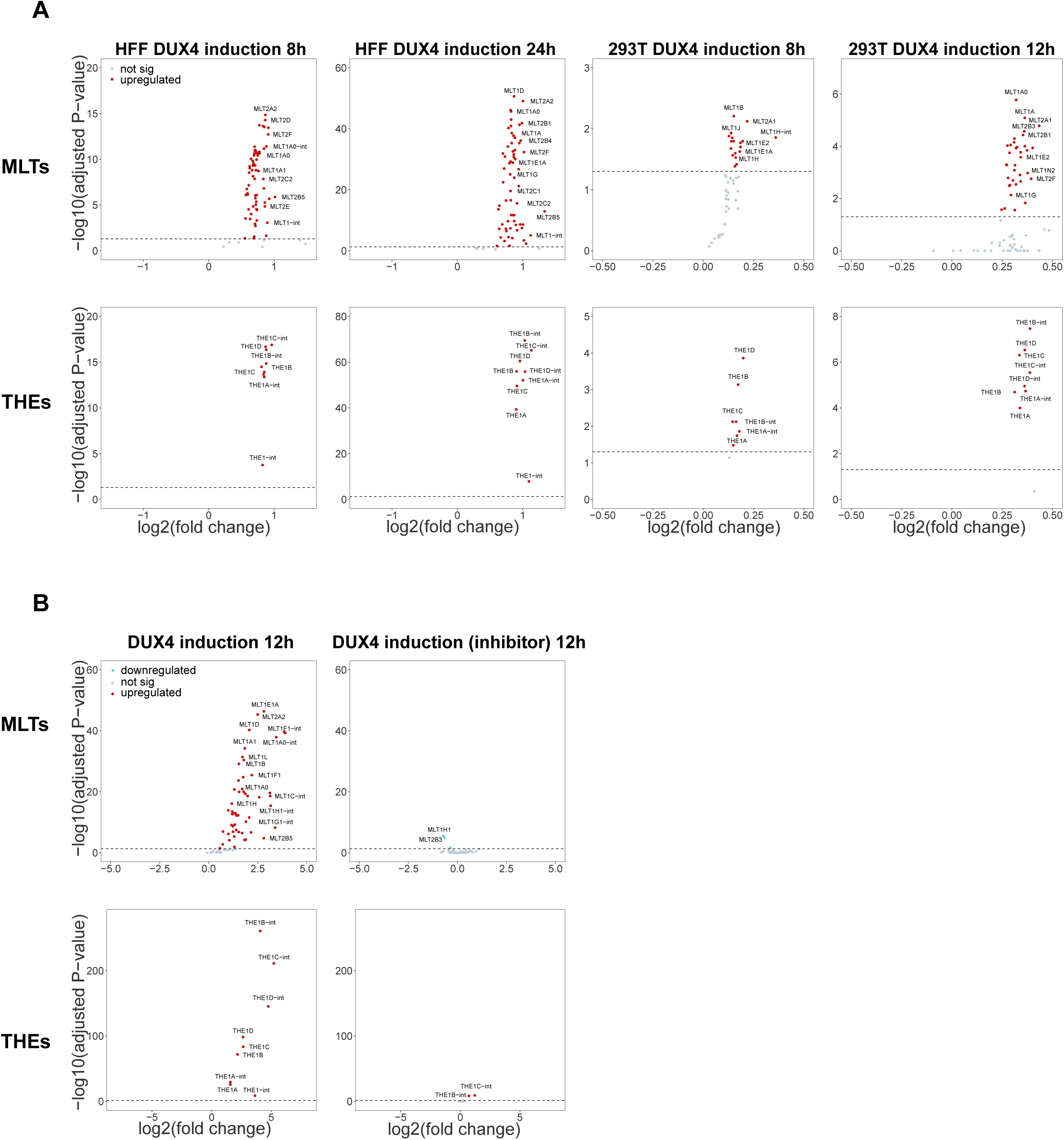
(**A**) Volcano plots showing significantly upregulated TEs of the MLT- and THE-families upon DUX4 induction for 8 and 24 hours in HFF cells and 8 and 12 hours in 293T cells. Red dots indicate TEs meeting the criteria of log2(FC) > 0 and padj < 0.05. (**B**) Volcano plot showing significantly upregulated MLTs and THEs upon DUX4 induced at 12 hours in myoblasts cells without and with DUX4 inhibitor treatment. Red dots indicate TEs meeting the criteria of log2(FC) > 0 and padj < 0.05.

**Fig. S15.**
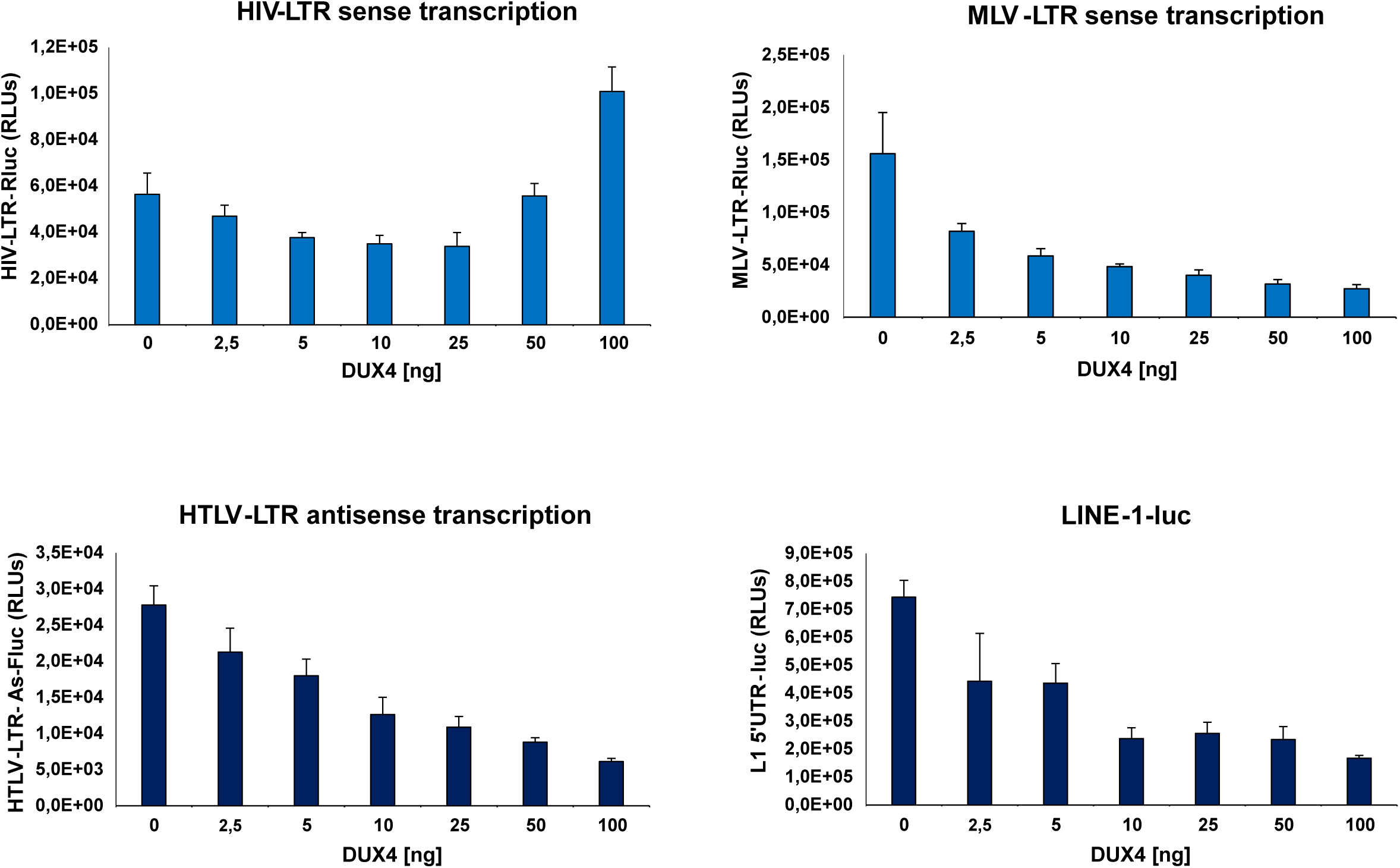
DUX4 affects HIV-LTR, MLV-LTR, HTLV-LTR and LINE-1 transcription. The graphs show the average RLUs of Luciferase assays of LTR-Luciferase constructs in the presence and absence of DUX4 of one representative of n=3 experiments. Error bars represent the standard deviation of the mean.

**Fig. S16.**
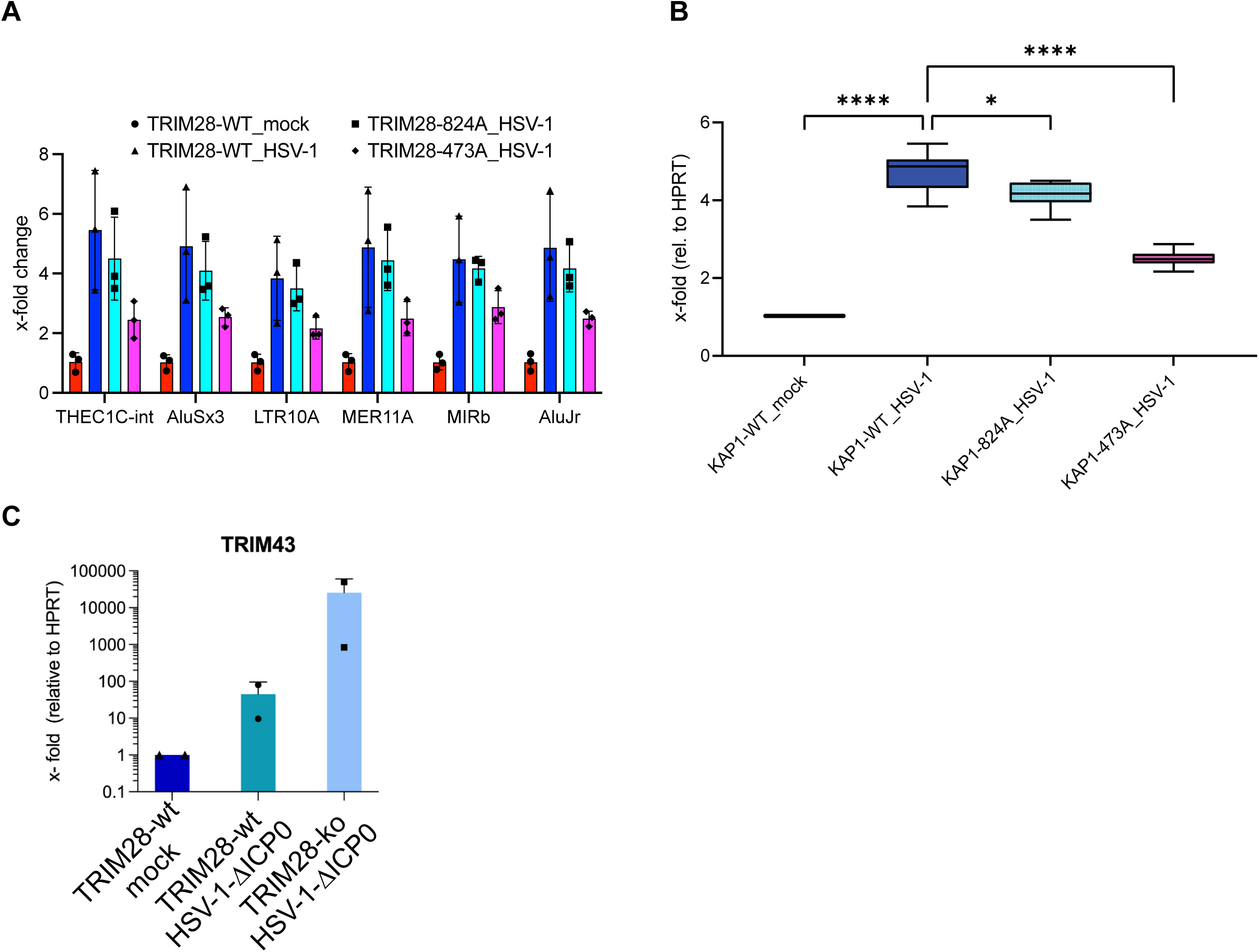
(**A**) Bar plot showing the fold change of TE-expression in A549 KAP1-WT cells (KAP1-KO rescue with reconstitution of WT KAP1), KAP1 -824A (KAP1-KO rescue with reconstitution of KAP1 S824A) and KAP1-473A (KAP1-KO rescue with KAP1 S473A) upon HSV-1 infection for 8 hours (hpi) by RT-qPCR. The 6 TEs MER11A, LTR10A, THE1C-int, AluSx3, AluJr and MIRb were upregulated upon HSV-1 infection in the RNAseq data. (**B**) Box plot showing the fold change of total 6 TEs expression in A549 KAP1-WT cells, KAP1 -824A and KAP1-473A upon HSV-1 infection for 8 hours (hpi) by RT-qPCR. Statistical significance was determined using two-sided nonparametric Wilcoxon rank-sum tests (ns: not significant, *P < 0.05, **P < 0.01, ***P < 0.001, ****P < 0.0001). (**C**) Bar plot showing the fold change of TRIM43 expression relative to HPRT in A549 KAP1-WT cells (KAP1-KO rescue with reconstitution of WT KAP1) and KAP1-KO cells upon HSV-1 ΔICP0 infection for 8 hours (hpi) by RT-qPCR.

**Fig. S17.**
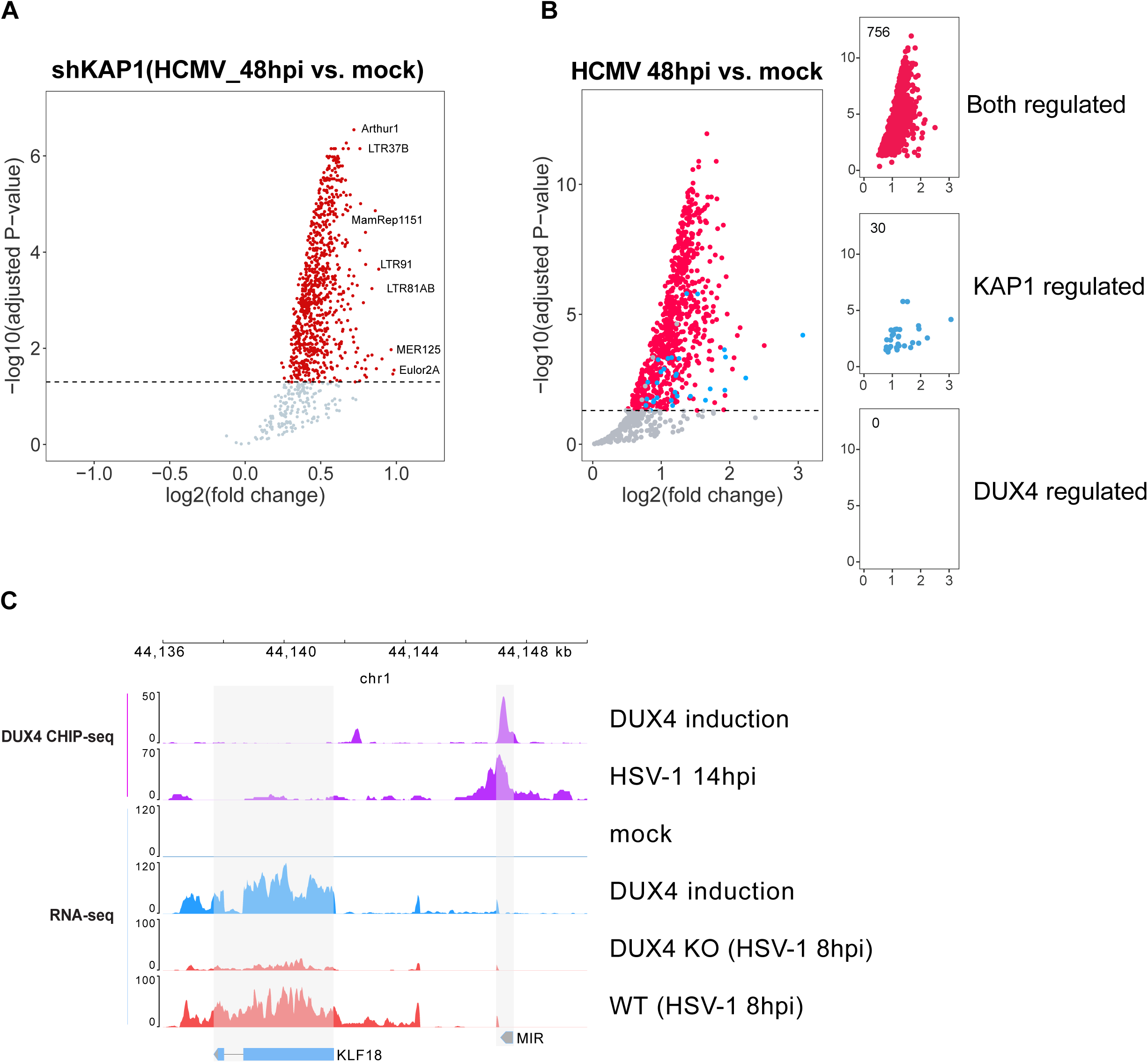
(**A**) Volcano plot illustrating upregulated TEs in HFF cells transfected with shKAP1 upon HCMV infection for 48 hpi. Red dots represent significantly upregulated TEs from the TEs analysis (padj < 0.05). (**B**) Volcano plot showing upregulated TEs upon HCMV (TB40/E) infection at 48 hpi that overlapped with upregulated Tes upon DUX4 induction (for 24 hours) and TEs regulated upon KAP1 knockdown in HFF cells. Red dots represent TEs significantly regulated by both KAP1 and DUX4, blue dots represent TEs regulated by KAP1, and purple dots represent TEs upregulated by DUX4 (log2(FC)>0, padj < 0.05). (**C**) Schematic representation and CHIP-seq tracks illustrate the binding of DUX4 to the TEs MIR that is located in the promoter region of the gene KLF18 after DUX4 induction and HSV-1 infection.

**Fig. S18.**
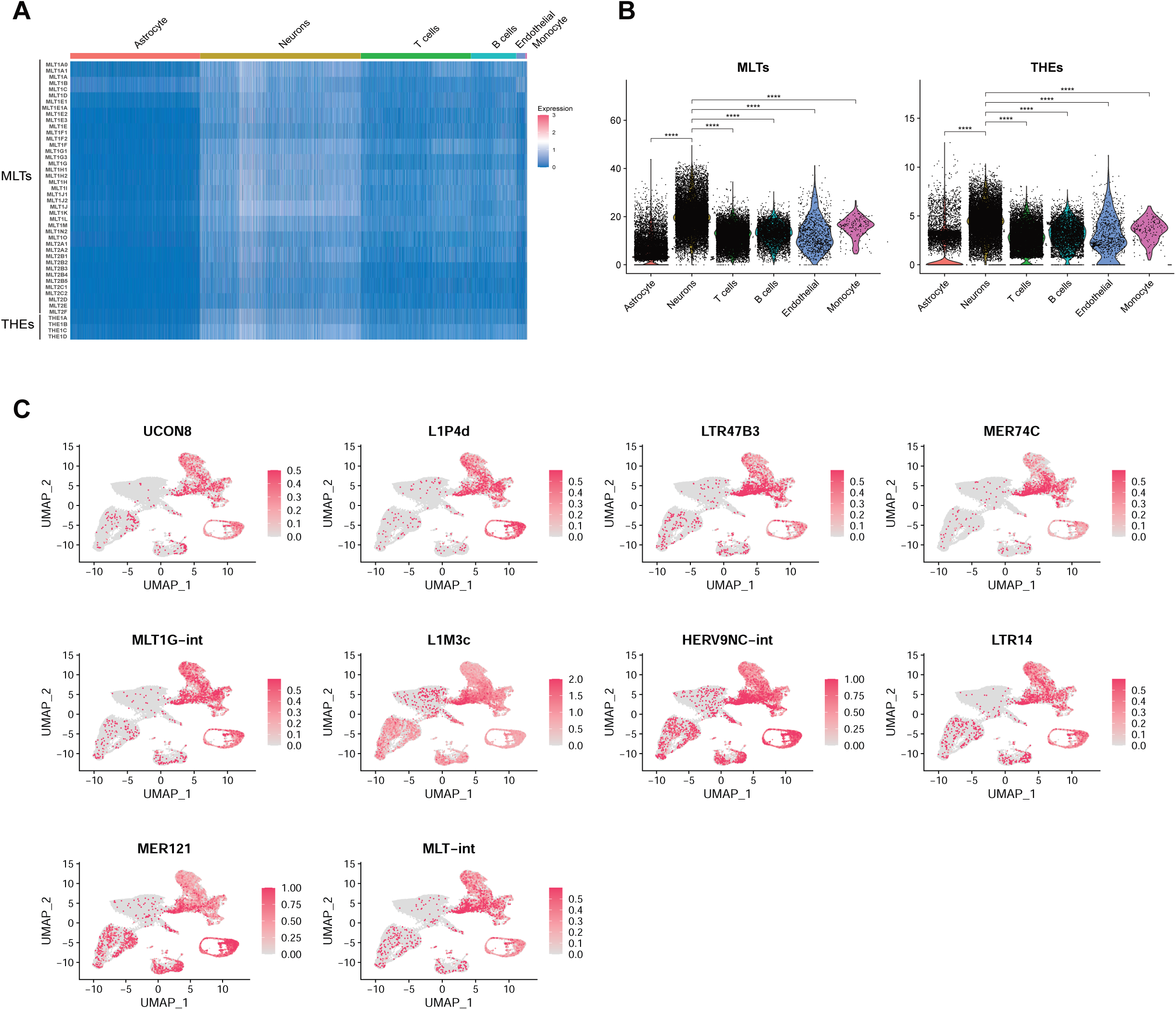
Single-Cell RNA-seq Analysis of TEs in Merkel Cell Carcinoma (MCC) tumor patients. **(A)** The heatmap illustrates the normalized read numbers of MLT and THE members across all cell types in Merkel Cell Carcinoma (MCC) tumor patients positive for Merkel Cell Polyomavirus (MCV). **(B)** MCV positive Merkel Cell Carcinoma (MCC) tumor patients: A series of violin plots that highlight the distribution of the normalized MLTs reads per cell and normalized THEs reads per cell in different cell types respectively. Statistical significance was determined using two-sided nonparametric Wilcoxon rank-sum tests (*P < 0.05, **P < 0.01, ***P < 0.001, ****P < 0.0001). **(C)** Examples of TEs enriched in cells with MCV and DUX4 target genes, more TEs examples are in the supplemental excel file.

**Fig. S19.**
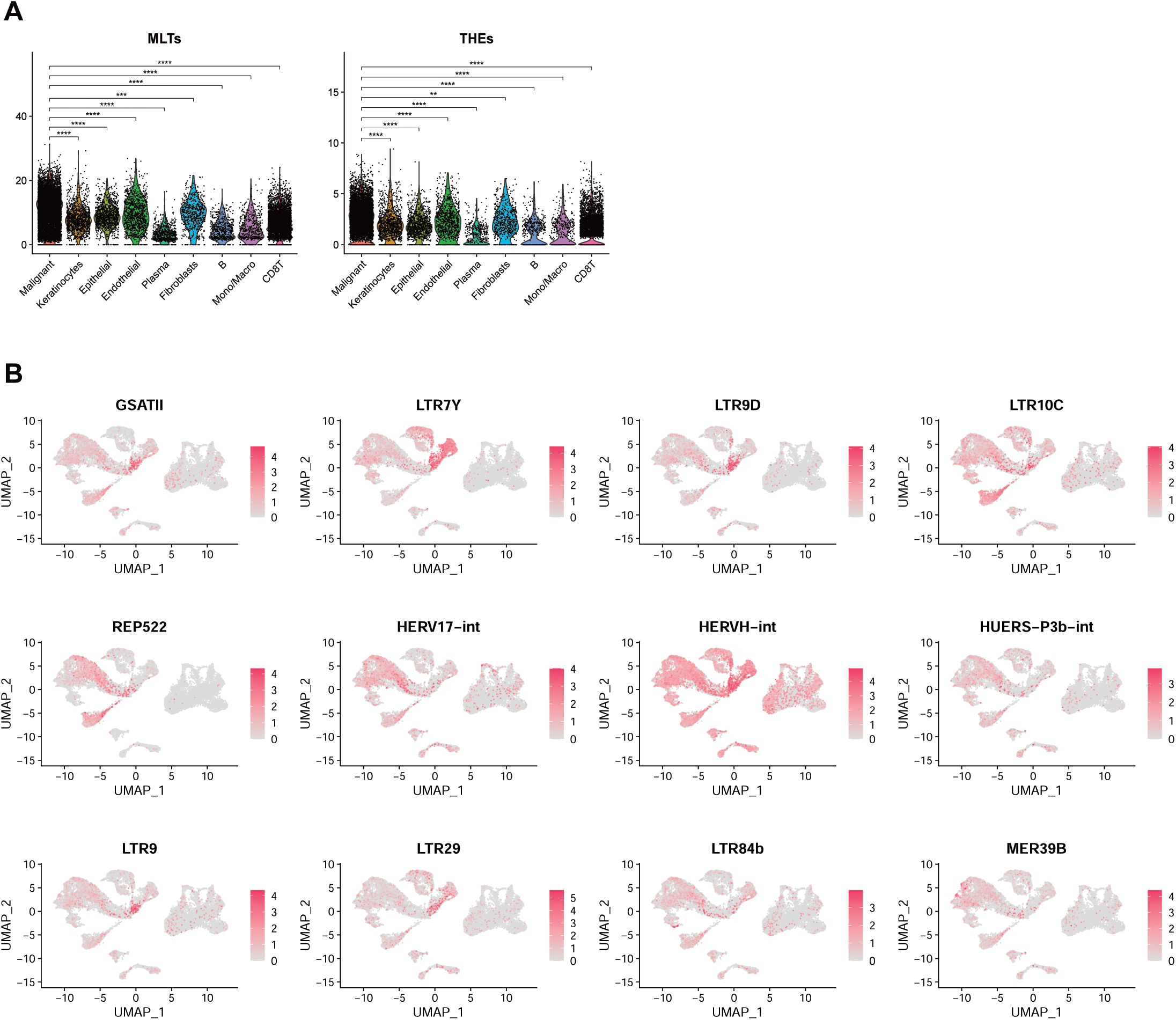
Single-Cell RNA-seq Analysis of TEs in Head and Neck Squamous Cell Carcinoma (HNSCC) Patients. **(A)** Head and Neck Squamous Cell Carcinoma (HNSCC) Patients with Human Papillomavirus (HPV) Positive: A series of violin plots that highlight the distribution of the normalized MLTs reads per cell and normalized THEs reads per cell in different cell types respectively. Statistical significance was determined using two-sided nonparametric Wilcoxon rank-sum tests (*P < 0.05, **P < 0.01, ***P < 0.001, ****P < 0.0001). **(B)** The examples of TEs enriched in cells with HPV and DUX4 target genes, more TEs examples are in the supplemental excel file.

## Notes

### Competing Interest Statement

The authors have declared no competing interest.

### Summary of Updates

The updated version of the manuscript includes several new figures

